# Lipid Landscape of the Human Retina and Supporting Tissues Revealed by High Resolution Imaging Mass Spectrometry

**DOI:** 10.1101/2020.04.08.029538

**Authors:** David M.G. Anderson, Jeffrey D. Messinger, Nathan H. Patterson, Emilio S. Rivera, Ankita Kotnala, Jeffrey M. Spraggins, Richard M. Caprioli, Christine A. Curcio, Kevin L. Schey

**Author notes:** Corresponding Author: Kevin L. Schey, Mass Spectrometry Research Center, PMB 407916, Nashville, TN 37240-7916.

## Abstract

The human retina evolved to facilitate complex visual tasks. It supports vision at light levels ranging from starlight to sunlight, and its supporting tissues and vasculature regulate plasma-delivered lipophilic essentials for vision, including retinoids (vitamin A derivatives). The human retina is of particular interest because of its unique anatomic specializations for high-acuity and color vision that are also vulnerable to prevalent blinding diseases. The retina’s exquisite cellular architecture is composed of numerous cell types that are aligned horizontally, giving rise to structurally distinct cell, synaptic, and vascular layers that are visible in histology and in diagnostic clinical imaging. Suitable for retinal investigations, MALDI imaging mass spectrometry (IMS) technologies are now capable of providing images at low micrometer spatial resolution with high levels of chemical specificity. In this study, a multimodal imaging approach combined with a recently developed method of high accuracy multi-image registration was used to define the localization of lipids in human retina tissue at laminar, cellular, and sub-cellular levels. Data acquired by IMS combined with autofluorescence and bright-field microscopy of human retina sections in macular and peripheral regions indicate differences in distributions and abundances of lipid species across and within single cell types. Of note is localization of signals within specific layers of macula, localization within different compartments of photoreceptors and RPE, complementarity of signals between macular retina and non-macular RPE, and evidence that lipids differing by a single double bond can have markedly different distributions.

## Introduction

The human retina is comprised of many specialized cell types including five types of neurons: photoreceptors, bipolar cells, ganglion cells, horizontal cells, and amacrine cells^1^ that are collectively responsible for photon absorption, signal transduction, and signal transmission to the visual centers of the brain. In addition, the retina contains cells that support neurons including Müller (radial) glia, retina pigment epithelial (RPE) cells, and vascular endothelium. These cells are horizontally aligned in an exquisite architecture, giving rise to structurally distinct layers of cell bodies, synaptic endings, and vascular beds that are visible in histology and in diagnostic clinical imaging. The human retina is of particular interest, because of its unique anatomic specializations for high-acuity and color vision^2^ that also renders it vulnerable to prevalent blinding diseases. Although the cellular architecture of human retina is well-characterized, its molecular composition is only partially understood. Complete knowledge of the molecular landscape of human retina is essential for understanding normal biochemistry and physiology as well as for understanding cellular dysfunction and pathology in disease.

Research over decades has highlighted the importance of lipids for retinal function. The content of polyunsaturated long chain fatty acids vary with cell type, age, and ocular pathologies^3-4^, and several monogenic inherited retinopathies directly impact lipid homeostasis and transfer^5-6^. The characteristic extracellular deposits (drusen) of age-related macular degeneration (AMD) are dominated by lipoprotein-related lipids of intraocular and dietary origin^7-9^. Retinoids (vitamin A derivatives) are essential for phototransduction and additionally feed into pathways resulting in amphiphilic fluorophores (bisretinoids) that are detectable in vivo and thus valuable for clinical autofluorescence imaging^10^. Molecular analysis; however, is challenging because the neurons are 1/10 the size of those in the brain and layers and spaces that are clinically relevant can be <10 μm thick^1, 11^.

Matrix assisted laser desorption ionization imaging mass spectrometry (MALDI IMS)^12^ provides the ability to map hundreds-to-thousands of molecular distributions in tissues at cellular resolution. With accurate co-registration to microscopy, these distributions can be correlated to very small histological features. MALDI IMS technology is particularly well-suited for examining tissues of the eye as demonstrated by published images of the retina^13-21^, optic nerve^14, 22-26^,lens^27-31^, and cornea^32^. Cell layers of the retina distinguished by IMS have unique layer-specific lipid and metabolite signatures^13-14, 16, 20^. With the development of multimodal imaging technologies incorporating data-rich IMS images with high spatial resolution methods like microscopy^33-36^, cellular and subcellular localization of specific molecules provide a powerful tool to understand biochemistry *in situ*. In this study, we used a recently developed method of high-accuracy registration^33^ to co-register high spatial resolution IMS images with histological images of the same tissue with single IMS pixel accuracy. This approach facilitated molecular characterization of aged healthy retinas to elucidate the molecular species in specific layers and cell types of central and peripheral human retina.

## Methods

### Tissue acquisition and characterization

Whole eyes were obtained from deceased human donors via Advancing Sight Network (Birmingham AL; formerly the Alabama Eye Bank) as part of studies on age-related macular degeneration (AMD). Donor criteria were ≥ 80 years of age, white, non-diabetic, and ≤6 hours death-to-preservation. In this demographic, AMD is prevalent, and eyes were screened for AMD presence and staging using *ex vivo* multimodal imaging including optical coherence tomography (OCT), a widely used clinical diagnostic imaging technology. Forty percent of eyes lacked detectable chorioretinal pathology and were available for the studies described herein.

### Tissue handling and *ex vivo* imaging

Methods were optimized for multimodal *ex vivo* clinical imaging of the ocular fundus^37^, i.e., the retina, choroid, and sclera. To create a consistent opening with a smooth contour, globes that were delivered intact by eye bank personnel, were opened in-house as follows. To facilitate removal of the cornea and 2-mm-wide scleral rim, a circular cut was scored with an 18-mm trephine (Stratis Healthcare, #6718L) and completed with a curved spring scissors (Roboz Cat# RS-5681). To facilitate preservative penetration without disturbing the vitreous body and detaching the retina from RPE, the iris was slit. Globes with lens and iris in place were immersed in buffered 4% paraformaldehyde overnight. The iris and lens were removed before imaging.

For imaging with OCT and scanning laser ophthalmoscopy, globes were immersed in buffer facing frontwards within a custom-built chamber with a 60-diopter lens ^37^. Spectral domain OCT (30° macula cube, 30 μm spacing, Automatic Real-time averaging ≥50), near-infrared reflectance, and autofluorescence (488 nm and 787 nm excitation wavelengths) images were captured with a Spectralis (HRA&OCT, HRA2; Heidelberg Engineering). For color fundus photographs, globes were placed within a 30 cc quartz crucible (Fisher Scientific # 08-074D). Images were taken at 3 magnifications (0.75x, 1.50x, 3.00x), and 3 lighting conditions (epi, flash, and dark field) with a Nikon D7200 camera mounted to an Olympus SZX9 Stereo microscope.

The goal of this study was to analyze macula and periphery together in continuous sections, since the two regions differ substantially by tissue mass, neuronal circuitry, and gene expression^38-40^. With respect to embedding in carboxymethylcellulose (CMC) prior to cryosectioning, the posterior pole was trimmed to a 14-mm-wide belt of retina, choroid, and sclera containing major landmarks (optic nerve head, fovea, and horizontal meridian of the visuotopic map) and extending anteriorly to pigmented tissue (ora serrata) at the edge of the ciliary body. To stabilize the globe and standardize dissection, posterior poles were placed in an custom-designed, chilled aluminum 3” x 4” x 1” billet with a 30-mm-diameter hemispheric well and a set of parallel grooves 7 mm superior and inferior to the center of the well (Supplemental Figure 1). Globes were placed facing up in the well and the globe-billet combination oriented so the superior quadrant appeared at the left and the grooves were vertical, thus making the long edge of the tissue block parallel to the optic nerve head and fovea axis. To guide a knife cut, the globe was snipped vertically (10 mm) with scissors (Roboz # 500216-G), guided by the parallel grooves. A tissue slicer (Thomas Sciences #6727C18) was placed in the two superior scissor cuts. With a guillotine motion, a superior cap was removed from the globe. While leaving the first blade in place to stabilize the globe, the inferior cap was removed by the same process at the second groove. The nasal periphery was removed with a slice 2 mm nasal to the optic nerve head.

Tissues were embedded in 2.5% carboxymethyl cellulose (Sigma C9481). Carboxymethyl cellulose powder was mixed in deionized water, heated to 70°C, stirred until dissolved (30 min -1 hr), and de-gassed before use. Tissue belts were placed into cryomolds (Peel-A-Way 22 x 30 mm molds, Polysciences # 18646B) using a stereo microscope, oriented so the superior edge faced up (to be sectioned first). Molds were filled with 5 ml of cold carboxymethyl cellulose and frozen at −20°C. For histopathologic evaluation, serial 10 µm cryosections were collected starting at the superior edge of the optic nerve head on pre-labeled 1×3 mm glass slides coated with 10% poly-L-lysine (Sigma Aldrich, St. Louis, MO, USA) and maintained at 37° during sectioning. At pre-defined intervals in the glass slide series, sections were captured on large, 45×45 mm in-house, poly-lysine coated indium-tin-oxide (ITO) slides (Delta Technologies Loveland, CO, USA) for IMS analysis. A slide box was vacuum-packed with an oxygen-absorbing packet within a Bitran freezer bag to help prevent oxidation and deterioration of lipid signal and stored at −80°C for transport to Vanderbilt.

### MALDI IMS Analysis

The matrices 2,5-dihydroxyacetophenone (DHA) and 1,5-diaminonaphalene (DAN) (Sigma Aldrich, St. Louis, MO, USA) were applied to tissue sections using a sublimation device developed in-house for positive and negative ion mode analysis, respectively. MALDI IMS data were acquired with a 10-15 µm pixel size in full scan mode using a Bruker SolariX 9.4T FT ICR mass spectrometer (Bruker Daltonics Billerica, MA, USA) equipped with a modified source with a Gaussian profile Nd:YAG laser and optics to provide a laser spot diameter of 8 µm. Following data acquisition, an advanced image registration workflow^33^ was performed using both autofluorescence and bright-field microscopy images. IMS data were exported for accurate registration and overlaid images were reconstructed using ImageJ Fiji freeware (NIH) while unregistered MALDI IMS images were generated using flexImaging (Bruker Daltonics, Billerica, MA, USA). Data were acquired with 500 shots per pixel and a mass range of *m/z* 154-2000 using the 1M data size resulting in a 0.5592 second transient, mass resolution was ∼60,000 at *m/z* 699.499.

### Microscopy

Pre-IMS autofluorescence (AF) images were acquired prior to matrix application on a Nikon Eclipse 90i (Nikon Instruments Inc, Melville, NY, USA) with a 10x objective (numerical aperture 0.30, 0.92 pixel/µm) using filter cubes for DAPI (excitation 325-375 nm, emission 435-485 nm) and Texas Red (excitation 540-580 nm, emission 600-660 nm). To record the laser burn pattern and assess tissue morphology, post-IMS AF images were taken following MALDI IMS using only the DAPI filter and low-intensity transmitted light. Following post-IMS AF microscopy, the matrix was removed using a minimal amount of methanol until the slide was visibly clear of matrix. The tissue was stained with hematoxylin (Mayer’s, MHS32-1L, Sigma Aldrich) and eosin (E6003, Sigma Aldrich) (H&E), and bright-field images were acquired at 10x on the same microscope.

### Registration

Registration of the IMS data to microscopy was performed using an explicit IMS pixel-to-laser ablation mark registration method described previously ^33^, provided in the *MSRC Registration Toolbox* (https://github.com/nhpatterson/regToolboxMSRC), and outlined in Figure 1. Briefly, the post-IMS AF ablation mark microscopy image was used to link individual ablation marks to the theoretical pixel position image generated from the IMS metadata (x,y coordinates). Then, automated computational registration of H&E and pre-IMS AF images to the previously IMS registered post-IMS AF ablation mark image was performed using *MSRC Registration Toolbox* software. Finally, the three microscopy images (H&E, pre-IMS AF and post-IMS AF) were registered directly to the IMS data via the ablation mark pattern.

**Figure 1.**
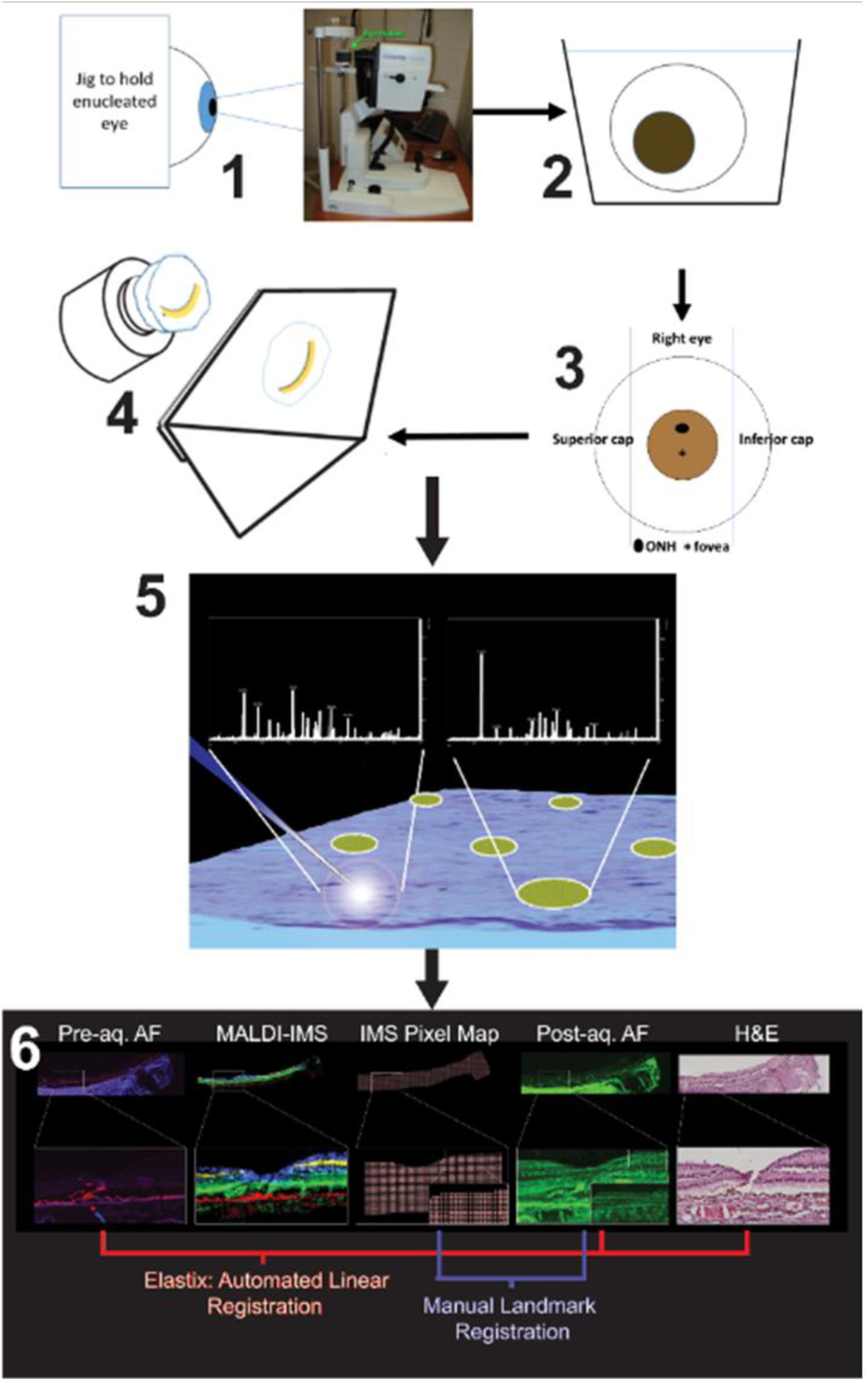
Workflow of sample preparation and multimodal imaging. **1.** Whole eye mounted in jig to obtain OCT image. **2.** The cornea is removed before fixation with 4% paraformaldehyde for 48 hours at 4° C. **3.** The eye is place in a dissection guide as described in the text to capture a belt with the optic nerve, macula, and temporal periphery. The belt is embedded in 2.6% CMC in a cryomold. The caps are discarded. **4.** Cryosections at 12-14 µm throughout the entire eye are thaw-mounted on either glass or ITO slides. **5.** ITO slides are imaged for autofluorescence (AF) before being coated with matrix via sublimation for acquisition of IMS data. **6.** Highly accurate data registration is performed from IMS, post-acquisition AF, pre AF, and H&E stained tissue.

### LC-MS Methods

Data acquisition was performed using a Vanquish UHPLC (Thermo Fisher Scientific, San Jose, CA) interfaced to a Q Exactive HF high resolution Orbitrap mass spectrometer (Thermo Fisher Scientific, San Jose, CA). Chromatographic separation was accomplished with a reverse phase Waters BEH C18 column, 2.1 mm x 150 mm, 1.7 µm (Waters, Milford, MA, USA) at a flow rate of 300 µL/min. The mobile phases were 10 mM ammonium acetate in A) water/acetonitrile (1:1), and B) acetonitrile/isopropanol (1:1). Ten µL of sample extracts were injected, and the elution gradient was as follows. The gradient was held at 20%B for the first minute, increased from 20%B to 100%B in 9 minutes where it was held for 30 s, and re-equilibrated at 20%B for 4.5 minutes. For all data acquired, spray voltage was set to 5 kV in negative mode with an S-lens RF level of 80 V, and capillary and probe temperatures of 320°C and 200°C, respectively. Sheath and auxiliary gases were set to 40 and 10 arbitrary units. Two different methods were used for collecting data in both untargeted and targeted modes:

#### Untargeted

Mass spectra were collected over a scan range of m/z 200 – 1500 at a resolution of 45,000. MS/MS spectra were collected using a top-7 data-dependent MS/MS method conducted at a resolution of 15,000, with an AGC target of 1e^5^, a max inject time of 50 ms, and a normalized collision energy of 30.

#### Targeted

Mass spectra were collected over a scan range of 500 – 1600 m/z, at a resolution of 30,000. MS/MS data were collected using parallel reaction monitoring at a resolution of 15,000 to isolate and fragment several ions of interest (see Supplemental Table 1). An isolation window of 2 m/z was used with a normalized collision energy of 30, AGC target of 2e^5^, and a maximum injection time of 100 ms.

#### Tentative molecular assignments

Assignments were made based solely on accurate mass measurements from IMS data when these had a <5 ppm error when fragmentation data were not obtained or were not interpretable.

## Results and Discussion

As the capabilities of IMS approach single cell or subcellular resolution, careful sample preparation and image registration between imaging modalities is required to maintain high quality tissue morphology and to assign exact molecular locations, respectively.

The preservation of native tissue morphology is paramount for obtaining high quality MALDI IMS images. Thus, the workflow utilized for sample preparation and image registration was crucial for localizing molecules to layers of the neural retina. Figure 1 describes the workflow from eye preservation to multimodal imaging performed in this study. The fixation and embedding protocols preserve tissue morphology in a manner that allows a MALDI process without suppression or contamination. In an autofluorescence image acquired prior to matrix application, the RPE layer can be clearly visualized due to its content of autofluorescent lipofuscin (long-lasting intracellular inclusion bodies) ^15, 41^. The explicit alignment of an IMS pixel to its corresponding laser burn pattern in images of autofluorescence and H&E-stained histology allows confident localization of signals to 0.5-2 µm accuracy, below the raster pitch of the IMS acquisition. Again, this capability is critical for imaging the precisely layered retina and single layer of RPE cells where inaccurate registration may lead to spurious tissue assignment of molecular signals. Utilizing morphology-rich microscopy images enables accurate registration to be achieved, providing high confidence when assigning laminar distributions of analytes.

### Central and Peripheral Retina Signals

Figure 2A shows a schematic of major retinal cell types distributed across the layers. In this description, the directions “inner” and “outer” indicate towards the center and exterior of the eye, respectively. Rod and cone photoreceptors responsible for dim and bright light vision, respectively, are vertically compartmentalized and horizontally aligned at the outer surface of the neurosensory retina. These cells initiate phototransduction at the outer segments (OS). Mitochondria are numerous in the ellipsoid (outer) part of the inner segments. Photoreceptor cell bodies stacked in rows comprise the outer nuclear layer (ONL). In the outer plexiform layer (OPL), cones contact bipolar and horizontal cells (interneurons) at large synaptic terminals called pedicles, and rods, at spherules. These interneurons have cell bodies in the inner nuclear layer (INL), and in the inner plexiform layer (IPL), they contact processes of ganglion cells. Müller glia (support cells to neurons) span the retina from inner limiting to external limiting membrane and are intercalated among the neurons in every layer. External to the photoreceptors, the retinal pigment epithelium (RPE) forms a supporting layer that is a single cell thick. The retina has a dual circulation system, a retinal circulation within the neurosensory retina with a physiologic barrier like the blood-brain barrier, and the choroidal circulation, a vascular bed behind the RPE and part of the systemic circulation. Photoreceptors and RPE are served by the choroid, with photoreceptor terminals also supplied by a plexus of the retinal circulation.

**Figure 2.**
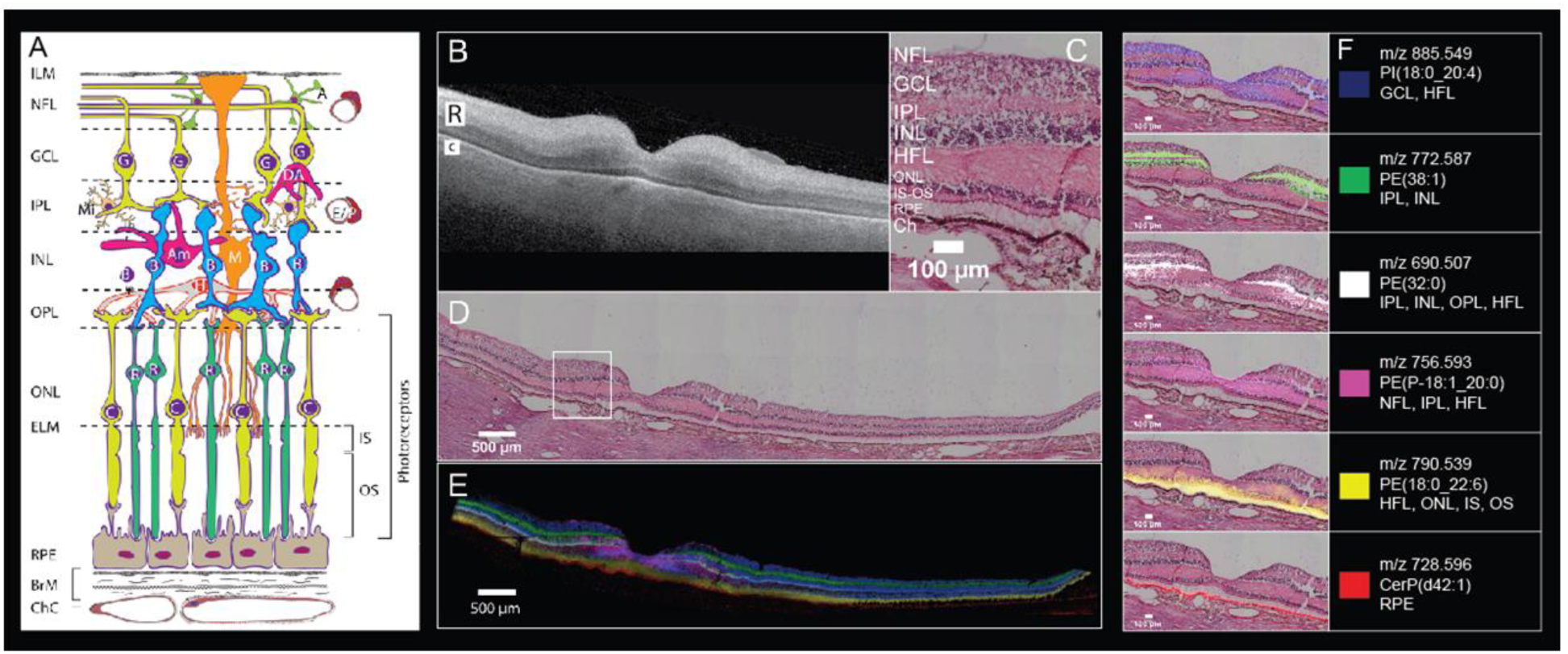
Six MALDI IMS signals with distinct laminar localizations in human macula from an 83-year-old female donor. **A)** Schematic diagram of retina, Bruch’s membrane and choriocapillaris, indicating cell types in their characteristic layers. The outer nuclear layer (ONL) contains cell bodies of cones (C) and rods (R) and their inner fibers, all interleaved with Müller glia (M). Photoreceptor inner segments (IS) contain many mitochondria. Photoreceptor outer segments (OS) contain phototransduction-related proteins in membrane disks. The RPE apical processes are specialized for metabolic exchange with photoreceptors and retinoid processing. Layers: ILM, inner limiting membrane; NFL nerve fiber layer, GCL ganglion cell layer, IPL inner plexiform layer, INL inner nuclear layer, HFL Henle fiber layer, ELM, external limiting membrane; RPE retinal pigment epithelium, ChC, choriocapillaris; BrM, Bruch’s membrane; cell types, A, astrocyte; Am, amacrine; B, bipolar; E, endothelium; G, ganglion cell; H, horizontal cell; P, pericyte. **B)** OCT B-scan of a donor eye macula prior to sample preparation, R and C indicating retina and choroid, respectively. The retina is at the top, the choroid (vasculature) is on the bottom, and the hyperreflective interface is the retinal pigment epithelium (RPE) layer. **C)** Zoomed image of tissue section stained with H&E to indicate cell layers at the level shown in A. NFL nerve fiber layer, GCL ganglion cell layer, IPL inner plexiform layer, INL inner nuclear layer, HFL Henle fiber layer, ONL outer nuclear layer, IS-OS photoreceptor inner and outer segments, RPE retinal pigment epithelium, Ch choroid. **D)** Panoramic H&E stained tissue post-IMS data acquisition showing macula and extended stretch of temporal peripheral retina. **E)** Overlay of MALDI IMS images of selected lipid signals localizing to distinct retinal layers and RPE as indicated. Colors defined in Panel F. **F)** Individual ion signals overlaid on top of H&E stained tissue image.

Figure 2B-E reveals retinal and choroidal structure in macular and peripheral regions of a normal retina of an 83-year-old donor. Figure 2B shows an optical coherence topography (OCT) B-scan of the preserved tissue before processing for IMS. Retinal layers are apparent despite of loss of transparency after death. The two most reflective bands in the neurosensory retina (Figure 2B “R”) are the IPL and OPL, and the RPE is a reflective band along the top (inner) edge of the choroid (Figure 2B “C”). The 6 mm diameter macula is defined by neuronal content and includes the fovea (1 mm diameter with a 0.35 mm, all-cone, rod-free zone in the middle), the parafovea (an annulus of 0.5-1.5 mm radius from the foveal center), and the surrounding perifovea (1.5-3 mm radius). The term “central macula” in this report refers to fovea and parafovea.

Figure 2C, D shows a panoramic (D) and zoomed (C) view of post-IMS H&E stained tissue, indicating the cell and synaptic layers of human macula and temporal periphery (Figure 2D). The distributions of six different molecular species are shown as an overlay (Figure 2E) and as individual ion images overlaid on the registered H&E-stained tissue image (Figure 2F). These figures demonstrate that the observed lipid species are highly localized to distinct layers. The PI(18:0_20:4) (m/z 885.549, blue) signal localizes to the GCL and INL as well as in the HFL and ONL. A signal observed at *m/z* 772.587 (green) and tentatively assigned as PE(38:1) is observed in the IPL and INL spanning from the parafovea region to the far periphery. The signal observed at *m/z* 756.593 (pink) was identified as a phosphatidylethanolamine plasmalogen (PE-P-18:1_20:0) and is localized to the central fovea and in the HFL, IPL and NFL extending only to the parafovea, unlike the other signals. The distribution observed for *m/z* 756.593 bears a striking resemblance to the ‘bowtie’ image of yellow xanthophyll carotenoid distribution, first described by Snodderly *et al.* using two-wavelength micro-densitometry of sectioned monkey retina ^42-45^. Because good evidence indicates that Müller glia in central macula are a major reservoir for xanthophylls ^46^, it is plausible that the *m/z* 756.593 bowtie may localize specifically to these cells.

A signal at *m/z* 690.507, tentatively identified as PE(32:0) (white), resides in the OPL, IPL, HFL, and INL, spanning the full breadth of the macula with lower abundance in the periphery. An intense signal at *m/z* 790.539 extending across the retina in the inner and outer segments (IS, OS) of photoreceptor (yellow) was identified as PE(18:0_22:6). This lipid is also observed in the HFL and ONL with lower abundance. This signal, previously observed in human photoreceptors by Zemski Berry *et al.* ^14^, is from a lipid that contains docosahexaenoic acid. This highly unsaturated fatty acid is abundant in neural tissue and especially in photoreceptors, where it is crucial for cellular function ^47-48^. External to the photoreceptors, a strong signal tentatively identified as a ceramide-1-phosphate (m/z 728.596, CerP(d42:1); red) is observed within the RPE layer across the entire retina (details below). Distributions of lipids in Figure 2 were replicated in the eye of an 81-year-old donor (Supplemental Figure 2). Lipid identification data can be found in Supplemental Table 1.

### Signals specific to photoreceptors and RPE

Rod photoreceptors are slender (2-3 µm) and can be very long (100 µm). They are the dominant photoreceptor in human retina (20:1 rod:cone ratio), being absent only in the cone-only foveal center. Signals unique to photoreceptor cells, a population necessarily dominated by rods in cross-sections not including the fovea, are shown in Figure 3.

**Figure 3.**
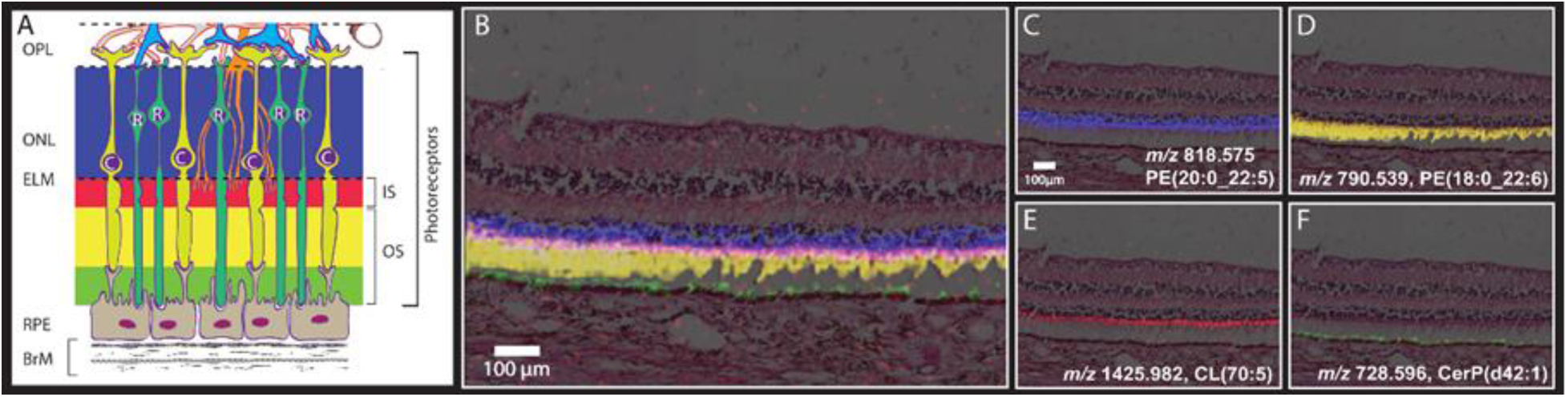
MALDI IMS signals consistent with localization to photoreceptor and RPE compartments. **A.** Schematic diagram of outer retina and Bruch’s membrane, excerpted from Figure 2A. Blue, pink, yellow, and green bands indicate layers formed by highly compartmentalized and vertically-aligned photoreceptors and RPE cells in panels B,C. See Figure 2 for explanation of cellular and subcellular content of each layer. Layers: OPL, outer plexiform layer; ONL, outer nuclear layer; ELM, external limiting membrane; RPE, retinal pigment epithelium; BrM, Bruch’s membrane; R, Rod; C, cone photoreceptors. **B-F.** Images and H&E stained tissue images overlaid in peripheral retina displaying signals from multiple lipid classes that localize to subcellular compartments of the photoreceptor cells. **B.** Overlay showing 4 separate signals defined in Panels C-F. **C.** Localized to ONL. **D.** Localized to photoreceptor inner and outer segments. **E.** Localized to mitochondria-rich photoreceptor inner segments. F. Localized to RPE apical processes.

Figure 3A shows part of the schematic from Figure 2A, focusing on photoreceptors and support cells. The RPE sends delicate processes in the apical direction to contact photoreceptor OS, near the RPE cell body for rods and 10-15 µm above the cell body for cones (because cone OS are short). The RPE sits on Bruch’s membrane, which serves as a flat vessel wall above the choriocapillaris endothelium. Figure 3A is color-coded to indicate photoreceptor and RPE compartments associated with IMS signals in Figure 3B (blue, red, yellow, green, for ONL, IS, OS, and RPE, respectively).

Figure 3B shows MALDI IMS images overlaid with an H&E image from this donor. The photoreceptors are attached to the RPE on the left of the image and detached from the RPE, a common artifact, on the right. A signal at *m/z* 818.575, identified as PE(20:0_22:6; blue), is observed with high abundance in the ONL, which contains photoreceptor cell bodies and processes of Müller glia. The *m/z* 818.575 signal was assigned to photoreceptors due to the lack of similar signal in other retinal layers where Müller glia are also present. Just external to this signal, a signal at *m/z* 1425.982 (red) can be seen in high abundance along a narrow band aligned with photoreceptor IS. This signal was identified as a cardiolipin CL(70:5). Cardiolipins are highly abundant in mitochondria, and the ellipsoid portion of photoreceptor IS contain many mitochondria providing energy for these metabolically active cells. This signal is also observed in inner retinal cells with abundant mitochondria. Below this layer are photoreceptor OS, where the docosahexaenoate-containing PE(18:0_22:6, Figure 2) is observed at *m/z* 790.539 (yellow). At the lowest part of the photoreceptor cells are OS that are interleaved with the apical processes of RPE cells. The signal observed at *m/z* 728.596 (green) is localized above and within the RPE, as discussed further below. Replicate data from an 81-year-old donor eye can be seen in Supplemental Figure 3.

The image of m/z 728.596 CerP(d42:1) highlights challenges posed by the precisely vertical compartmentalization of the RPE, a critical layer in initiating vision and also central to AMD pathogenesis. The RPE performs demanding dual services to photoreceptors above and choriocapillaris below, as reflected in an internal gradient of organelles and molecules. For example, delicate apical processes contacting photoreceptors (Figure 2A, 3) and cell bodies share retinoid processing enzymes but differ in channels important for potassium currents ^49-50^. Figure 2 and Supplemental Figure 4 overlaying IMS with tissue autofluorescence show strong m/z 728.596 signal in RPE. Upon closer examination of Figure 2 this signal appears to extend above the RPE layer by up to 10 µm, depending on retinal position, with the greatest extensions observed at the fovea. Conversely, Figure 3 and Supplemental Figure 5 show signal localized *only* to apical processes, i.e., not in the cell bodies, especially where apical processes are fortuitously standing upright. These discrepancies in locations are due to the different resolutions used (10 µm for Figures 2-5 vs 15 µm for Supplemental Figures 2-3) and whether or not OS are attached to RPE. These differences will be clarified by processing more samples at the higher resolution.

**Figure 4.**
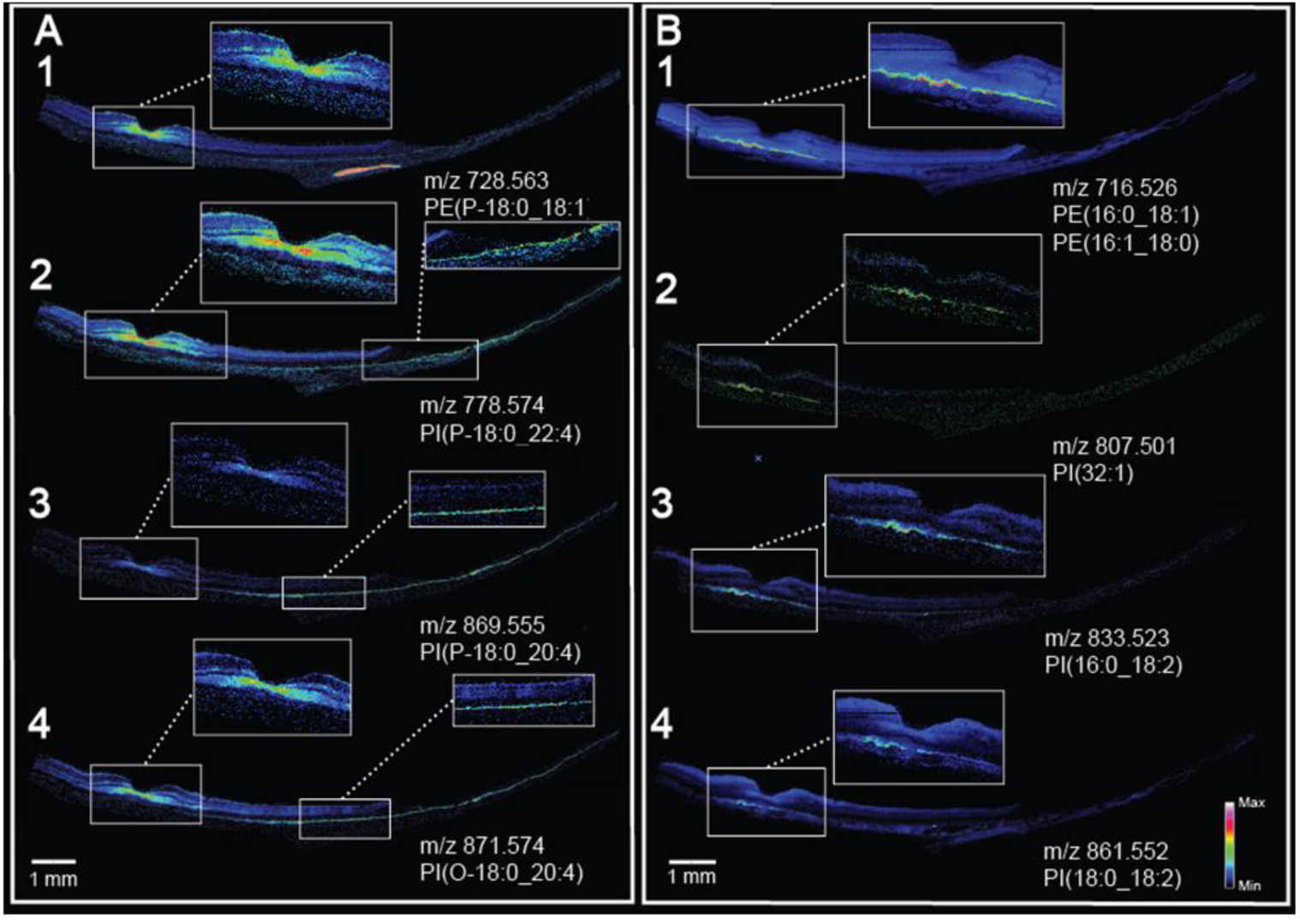
MALDI IMS shows complementarity of signals in retina and RPE. **A)** Signals localized to the foveal center and extending into the macular NFL, IPL, HFL also localize to the peripheral RPE. **B)** Signals localized to RPE underlying the macula, not confined to the fovea.

**Figure 5.**
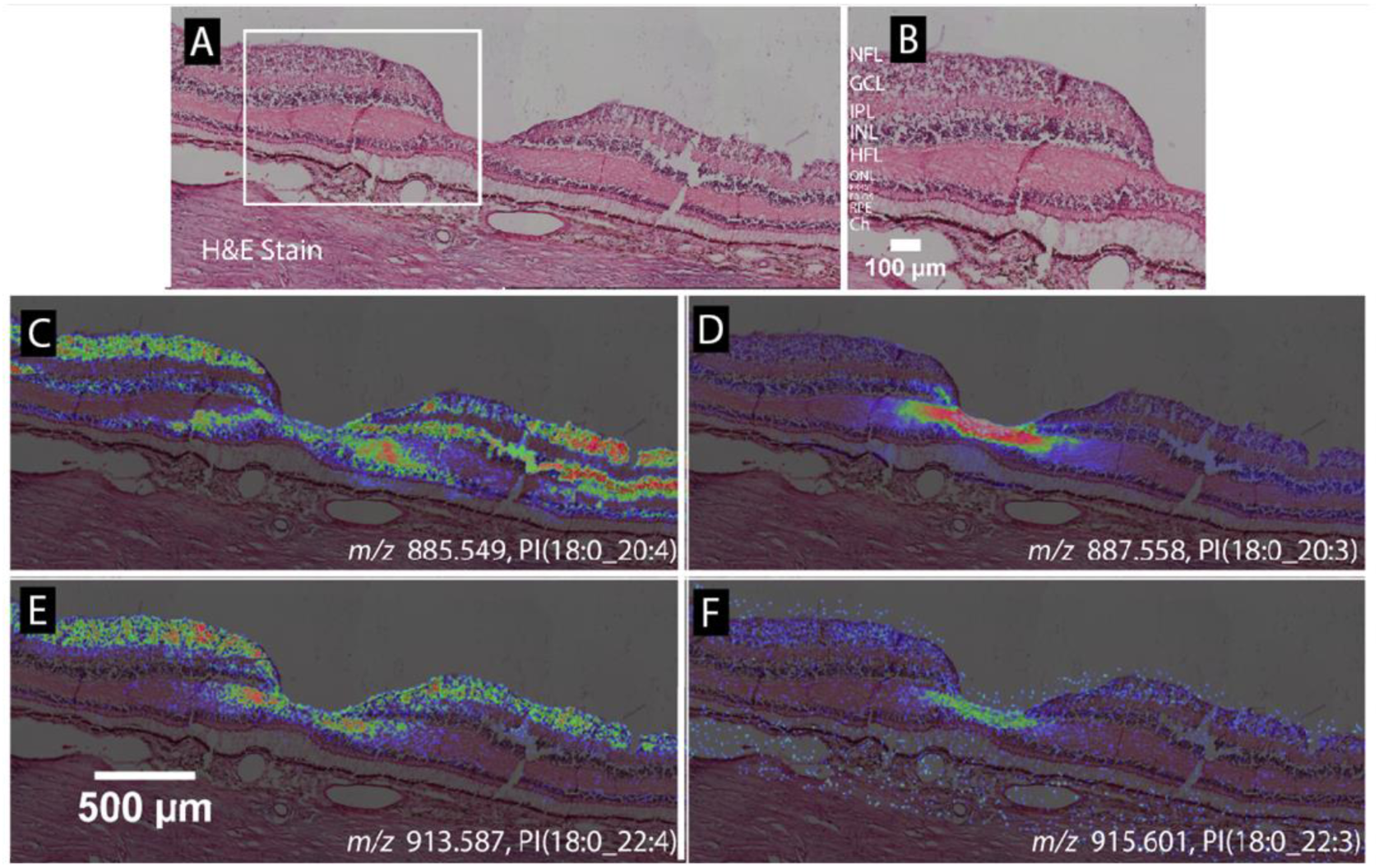
Lipid species varying in only one double bond can exhibit different MALDI IMS distributions. **A)** H&E stained tissue image of a normal macula, with the foveal pit in the center. **B)** Zoomed view of panel A showing retinal layers. NFL, nerve fiber layer; GCL, ganglion cell layer; IPL, inner plexiform layer; INL, inner nuclear layer; HFL, Henle fiber layer; ONL, outer nuclear layer; PR/IS, photoreceptor inner segments; PR/OS, photoreceptor outer segments; RPE, retinal pigment epithelium; ChC, choriocapillaris. **C)** *m/z* 885.5499, PI(18:0_20:4) distributes in the foveal center, GCL, INL, and HFL. **D)** *m/z* 887.558, PI(18:0_20:3), with one double bond less than the species in panel C, localizes to the HFL of the foveal center and the inner INL. **E)** *m/z* 913.587, PI(18:0_22:4) localizes to the foveal center, GCL, INL, and HFL, like the species in panel A, but with a lesser lateral extent of INL. **F)** *m/z* 915.601, PI(18:0_22:3), with one double bond less than the species in panel E, localizes to the HFL of the foveal center.

### Complementarity of neurosensory retina and RPE

Figure 4 displays eight MALDI IMS images from lipid species that exhibit complementary localization patterns in the macula of neurosensory retina and underlying RPE. Figure 4A shows images generated from four plasmalogen phosphatidylglycerol lipid species. Panels A1-4 all exhibit high signal in the foveal center. Panels A1, 2 and 4 also have strong signal in the HFL, NFL, and IPL extending across the whole macula. Figure 4A3 displays a similar overall pattern with high signal in the peripheral retina but lower signal in the central macula. In all 4 panels, no signal is observed in RPE directly below the signal-rich area of neurosensory retina, yet signals are intense in peripheral RPE.

Conversely, Figure 4B1-4 shows four signals that exhibit high abundance in the RPE in an area ∼3 mm wide and centered under the fovea, but not confined to the rod-free area, and exhibiting no signal in the periphery. The signal observed at *m/z* 716.526 in Figure 4B1 was attributed to two phosphatidylethanolamine lipids PE(16:0_18:1) and PE(16:1_18:0) that are exact isobars, as confirmed in LC-MS/MS experiments. The MALDI IMS image generated from three phosphatidylinositol lipids, one tentatively identified as PI(32:1) (Figure 4B2) and the other two identified as PI(16:0_18:2) and PI(18:0_18:2) (Figure 4B 3-4) all display the same distribution but with lower relative intensities than that observed for the PE lipid in Figure 4B1. The positively identified PI lipids contain an 18:2 fatty acid side chain. The same distributions were also observed in an 81-year-old donor eye as seen in Supplemental Figure 6. The relevance of the apparent high abundance and specific localization of this particular fatty acid side chain has yet to be determined, although variations in abundance between mid-life, aged normal, and AMD eyes was previously reported by Liu *et al* ^3^.

Overall, Figure 4 clearly indicates not only differences in molecular composition between central and peripheral RPE but also a regional complementarity of neurosensory retina and RPE. Further, this arrangement is not confined to the rod-free fovea but rather involves the central macula, a region distinguished by high concentration of xanthophyll carotenoids. Regarding results in Figure 4A, plasmalogen lipids can act as free radical scavengers and to be associated with glial cells, which may explain their localization in the foveal HFL along with cones. The role of ambient light is often invoked to explain free radical generation ^51-55^ in the retina. However, retinal illuminance in the human eye is nearly uniform for a wide portion of the visual field, and light distribution alone cannot explain the specific localizations observed in our studies ^56-57^. An alternative explanation for these localizations builds on the idea that in Müller glia in central macula have distinct properties, including xanthophyll sequestration and also, a separate retina-based retinoid regeneration cycle believed to serve cones ^58^. These pathways are distinct from the canonical visual cycle present in RPE ^59-60^.

We speculate that retina-RPE complementarity in Figure 4 indicates that similar functions are performed by macular Müller cells and extramacular RPE. Further, we hypothesize that these functions include retinoid processing, due to a division of labor between Müller cells and RPE unique to macula, as a result of a retinoid cycle serving cones. The localized lipids may or may not be directly involved in retinoid processing *per se.* They may signify an overall distinct internal milieu in the cells harboring these pathways. Finally, our data suggest that the molecular inhomogeneity of human RPE first seen for bisretinoid A2E (low signal in macula, high in periphery) ^15, 20^ is part of much larger pattern involving lipids in multiple essential functions in cells supporting neuronal function. The cellular and molecular basis for this retinal topography will be important to explore, because the layout of each species’ retina is evolved for efficient vision in its own habitat ^61^, under genetic control.

### Spectral specificity of high resolution IMS

Figure 5 demonstrates markedly different localizations for molecules differing by a single double bond. Figures 5A and B, respectively, show a panoramic and magnified image of an H&E stained macula with the fovea in the center. Figure 5C displays the distribution of PI(18:0_20:4) at m/z 885.549, localized to the foveal center, GCL, INL, and HFL. Figure 5D was generated from m/z 887.558, identified as PI(18:0_20:3) and differing by only one double bond from the lipid shown in Figure 5C, yet specifically localizing to the HFL in the fovea. Also localizing to these regions is another lipid pair varying in one double bond of a fatty acid chain, i.e., PI(18:0_22:4) and PI(18:0_22:3) shown in Figure 5E and F, respectively. Data shown in Figure 5 were replicated from an 81-year-old donor tissue as shown in Supplemental Figure 7. This observation of the same lipid with one extra double bond showing contrasting, and to some extent inverse, localization was recently observed in the human optic nerve ^23^. In this tissue, one species of sphingomyelin (SM(d36:2)) localized to axonal bundles while another (SM(d36:1)) was present in the surrounding glial cells and connective tissue ^23^.

## Conclusions

Registration of images from multimodal imaging experiments allows accurate assignment of IMS signals to retina layers, which, with knowledge of layer composition, can permit assignment to specific cell types and even subcellular compartments. The patterns shown in Figures 2-5 are only a subset of the many signals detected thus far. Supplemental Figures 8 and 9 display 100 images selected from the two donor tissues (83 and 81 year old) indicating the diverse localizations of the many signals observed.

This multimodal imaging method provides a powerful tool to elucidate the molecular composition of the numerous cell types in the neural retina and its supporting tissues. Use of *ex vivo* imaging with OCT allows clinically-relevant pathology to be identified prior to molecular analysis, a measure of quality control over post-mortem tissue destined for IMS, and translation of layer- and region-specific m/z signals to the interpretation of a widely used diagnostic technique for retinal disease. Data presented herein show the complexity of molecular distributions associated with the many cell types present in the neural retina and the striking topographies even within the same cell type. These data further demonstrate the regional heterogeneity of human RPE, of significance in understanding the RPE’s evolved role in supporting photoreceptors and vasculature and its dysfunction in macular disease. These data also set the stage for exploring the relevance of lipid localization and abundance in normal retinal function.

## Acknowledgements

This project was supported by the National Institutes of Health P41 GM103391 (RMC) and R01 EY027948 (CC).

## References

1. Masland, Richard H., The Neuronal Organization of the Retina. Neuron 2012, 76 (2), 266–280.

2. Rossi, E. A.; Roorda, A., The relationship between visual resolution and cone spacing in the human fovea. Nat Neurosci 2010, 13 (2), 156–7.

3. Liu, A.; Chang, J.; Lin, Y.; Shen, Z.; Bernstein, P. S., Long-chain and very long-chain polyunsaturated fatty acids in ocular aging and age-related macular degeneration. Journal of lipid research 2010, 51 (11), 3217–3229.

4. Agbaga, M.-P.; Merriman, D. K.; Brush, R. S.; Lydic, T. A.; Conley, S. M.; Naash, M. I.; Jackson, S.; Woods, A. S.; Reid, G. E.; Busik, J. V.; Anderson, R. E., Differential composition of DHA and very-long-chain PUFAs in rod and cone photoreceptors. Journal of Lipid Research 2018, 59 (9), 1586–1596.

5. Harkewicz, R.; Du, H.; Tong, Z.; Alkuraya, H.; Bedell, M.; Sun, W.; Wang, X.; Hsu, Y. H.; Esteve-Rudd, J.; Hughes, G.; Su, Z.; Zhang, M.; Lopes, V. S.; Molday, R. S.; Williams, D. S.; Dennis, E. A.; Zhang, K., Essential role of ELOVL4 protein in very long chain fatty acid synthesis and retinal function. J Biol Chem 2012, 287 (14), 11469–80.

6. Sluch, V. M.; Banks, A.; Li, H.; Crowley, M. A.; Davis, V.; Xiang, C.; Yang, J.; Demirs, J. T.; Vrouvlianis, J.; Leehy, B.; Hanks, S.; Hyman, A. M.; Aranda, J.; Chang, B.; Bigelow, C. E.; Rice, D. S., ADIPOR1 is essential for vision and its RPE expression is lost in the Mfrp(rd6) mouse. Sci Rep 2018, 8 (1), 14339.

7. Curcio, C. A., Soft Drusen in Age-Related Macular Degeneration: Biology and Targeting Via the Oil Spill Strategies. Invest Ophthalmol Vis Sci 2018, 59 (4), AMD160–AMD181.

8. Curcio, C. A., Antecedents of Soft Drusen, the Specific Deposits of Age-Related Macular Degeneration, in the Biology of Human MaculaWhy the Macula for Soft Drusen? Investigative Ophthalmology & Visual Science 2018, 59 (4), AMD182–AMD194.

9. Bretillon, L.; Thuret, G.; Gregoire, S.; Acar, N.; Joffre, C.; Bron, A. M.; Gain, P.; Creuzot-Garcher, C. P., Lipid and fatty acid profile of the retina, retinal pigment epithelium/choroid, and the lacrimal gland, and associations with adipose tissue fatty acids in human subjects. Exp Eye Res 2008, 87 (6), 521–8.

10. Sparrow, J. R.; Yoon, K. D.; Wu, Y.; Yamamoto, K., Interpretations of fundus autofluorescence from studies of the bisretinoids of the retina. Invest Ophthalmol Vis Sci 2010, 51 (9), 4351–7.

11. Curcio, C. A., Messinger, J. D., Sloan, K. R., McGwin, G., Jr, Medeiros, N. E., Spaide, R. F.,, Subretinal drusenoid deposits in non-neovascular age-related macular degeneration: morphology, prevalence, topography, and biogenesis model. Retina 2013, 33, 265–276.

12. Caprioli, R. M.; Farmer, T. B.; Gile, J., Molecular imaging of biological samples: localization of peptides and proteins using MALDI-TOF MS. Anal Chem 1997, 69 (23), 4751–60.

13. Anderson, D. M.; Ablonczy, Z.; Koutalos, Y.; Spraggins, J.; Crouch, R. K.; Caprioli, R. M.; Schey, K. L., High resolution MALDI imaging mass spectrometry of retinal tissue lipids. J Am Soc Mass Spectrom 2014, 25 (8), 1394–403.

14. Zemski Berry, K. A.; Gordon, W. C.; Murphy, R. C.; Bazan, N. G., Spatial organization of lipids in the human retina and optic nerve by MALDI imaging mass spectrometry. J Lipid Res 2014, 55 (3), 504–15.

15. Ablonczy, Z.; Higbee, D.; Anderson, D. M.; Dahrouj, M.; Grey, A. C.; Gutierrez, D.; Koutalos, Y.; Schey, K. L.; Hanneken, A.; Crouch, R. K., Lack of correlation between the spatial distribution of A2E and lipofuscin fluorescence in the human retinal pigment epithelium. Invest Ophthalmol Vis Sci 2013, 54 (8), 5535–42.

16. Roy, M. C.; Nakanishi, H.; Takahashi, K.; Nakanishi, S.; Kajihara, S.; Hayasaka, T.; Setou, M.; Ogawa, K.; Taguchi, R.; Naito, T., Salamander retina phospholipids and their localization by MALDI imaging mass spectrometry at cellular size resolution. J Lipid Res 2011, 52 (3), 463–70.

17. Garrett, T. J.; Dawson, W. W., Lipid geographical analysis of the primate macula by imaging mass spectrometry. Methods Mol Biol 2009, 579, 247–60.

18. Garrett, T. J.; Menger, R. F.; Dawson, W. W.; Yost, R. A., Lipid analysis of flat-mounted eye tissue by imaging mass spectrometry with identification of contaminants in preservation. Anal Bioanal Chem 2011, 401 (1), 103–13.

19. Palmer, A. D.; Griffiths, R.; Styles, I.; Claridge, E.; Calcagni, A.; Bunch, J., Sucrose cryo-protection facilitates imaging of whole eye sections by MALDI mass spectrometry. J Mass Spectrom 2012, 47 (2), 237–41.

20. Anderson, D. M. G.; Ablonczy, Z.; Koutalos, Y.; Hanneken, A. M.; Spraggins, J. M.; Calcutt, M. W.; Crouch, R. K.; Caprioli, R. M.; Schey, K. L., Bis(monoacylglycero)phosphate lipids in the retinal pigment epithelium implicate lysosomal/endosomal dysfunction in a model of Stargardt disease and human retinas. Scientific Reports 2017, 7 (1), 17352.

21. Bowrey, H. E.; Anderson, D. M.; Pallitto, P.; Gutierrez, D. B.; Fan, J.; Crouch, R. K.; Schey, K. L.; Ablonczy, Z., Imaging mass spectrometry of the visual system: Advancing the molecular understanding of retina degenerations. Proteomics. Clinical applications 2016, 10 (4), 391–402.

22. Anderson, D. M.; Mills, D.; Spraggins, J.; Lambert, W. S.; Calkins, D. J.; Schey, K. L., High-resolution matrix-assisted laser desorption ionization-imaging mass spectrometry of lipids in rodent optic nerve tissue. Mol Vis 2013, 19, 581–92.

23. Anderson, D. M.; Spraggins, J. M.; Rose, K. L.; Schey, K. L., High spatial resolution imaging mass spectrometry of human optic nerve lipids and proteins. J Am Soc Mass Spectrom 2015, 26 (6), 940–7.

24. Sato, K.; Saigusa, D.; Saito, R.; Fujioka, A.; Nakagawa, Y.; Nishiguchi, K. M.; Kokubun, T.; Motoike, I. N.; Maruyama, K.; Omodaka, K.; Shiga, Y.; Uruno, A.; Koshiba, S.; Yamamoto, M.; Nakazawa, T., Metabolomic changes in the mouse retina after optic nerve injury. Scientific Reports 2018, 8 (1), 11930.

25. Stark, D. T.; Anderson, D. M. G.; Kwong, J. M. K.; Patterson, N. H.; Schey, K. L.; Caprioli, R. M.; Caprioli, J., Optic Nerve Regeneration After Crush Remodels the Injury Site: Molecular Insights From Imaging Mass SpectrometryOptic Nerve Regeneration Imaging Mass Spectrometry. Investigative Ophthalmology & Visual Science 2018, 59 (1), 212–222.

26. Anderson, D. M. G.; Van de Plas, R.; Rose, K. L.; Hill, S.; Schey, K. L.; Solga, A. C.; Gutmann, D. H.; Caprioli, R. M., 3-D imaging mass spectrometry of protein distributions in mouse Neurofibromatosis 1 (NF1)-associated optic glioma. Journal of Proteomics 2016, 149, 77–84.

27. Grey, A. C.; Schey, K. L., Age-related changes in the spatial distribution of human lens alpha-crystallin products by MALDI imaging mass spectrometry. Invest Ophthalmol Vis Sci 2009, 50 (9), 4319–29.

28. Stella, D. R.; Floyd, K. A.; Grey, A. C.; Renfrow, M. B.; Schey, K. L.; Barnes, S., Tissue localization and solubilities of alphaA-crystallin and its numerous C-terminal truncation products in pre- and postcataractous ICR/f rat lenses. Invest Ophthalmol Vis Sci 2010, 51 (10), 5153–61.

29. Ellis, S. R.; Wu, C.; Deeley, J. M.; Zhu, X.; Truscott, R. J. W.; Panhuis, M. i. h.; Cooks, R. G.; Mitchell, T. W.; Blanksby, S. J., Imaging of Human Lens Lipids by Desorption Electrospray Ionization Mass Spectrometry. Journal of the American Society for Mass Spectrometry 2010, 21 (12), 2095–2104.

30. Pól, J.; Vidová, V.; Hyötyläinen, T.; Volný, M.; Novák, P.; Strohalm, M.; Kostiainen, R.; Havlíček, V.; Wiedmer, S. K.; Holopainen, J. M., Spatial distribution of glycerophospholipids in the ocular lens. PloS one 2011, 6 (4), e19441–e19441.

31. Le, C. H.; Han, J.; Borchers, C. H., Dithranol as a matrix for matrix assisted laser desorption/ionization imaging on a fourier transform ion cyclotron resonance mass spectrometer. Journal of visualized experiments : JoVE 2013, (81), e50733–e50733.

32. Chen, Y.; Jester, J. V.; Anderson, D. M.; Marchitti, S. A.; Schey, K. L.; Thompson, D. C.; Vasiliou, V., Corneal haze phenotype in Aldh3a1-null mice: In vivo confocal microscopy and tissue imaging mass spectrometry. Chemico-Biological Interactions 2017, 276, 9–14.

33. Patterson, N. H.; Tuck, M.; Van de Plas, R.; Caprioli, R. M., Advanced Registration and Analysis of MALDI Imaging Mass Spectrometry Measurements through Autofluorescence Microscopy. Analytical Chemistry 2018, 90 (21), 12395–12403.

34. Svirkova, A.; Turyanskaya, A.; Perneczky, L.; Streli, C.; Marchetti-Deschmann, M., Multimodal imaging of undecalcified tissue sections by MALDI MS and μXRF. Analyst 2018, 143 (11), 2587–2595.

35. Van de Plas, R.; Yang, J.; Spraggins, J.; Caprioli, R. M., Image fusion of mass spectrometry and microscopy: a multimodality paradigm for molecular tissue mapping. Nature methods 2015, 12 (4), 366–372.

36. Chughtai, S.; Chughtai, K.; Cillero-Pastor, B.; Kiss, A.; Agrawal, P.; MacAleese, L.; Heeren, R. M. A., A multimodal mass spectrometry imaging approach for the study of musculoskeletal tissues. International Journal of Mass Spectrometry 2012, 325-327, 150–160.

37. Pang, C. E.; Messinger, J. D.; Zanzottera, E. C.; Freund, K. B.; Curcio, C. A., The Onion Sign in Neovascular Age-Related Macular Degeneration Represents Cholesterol Crystals. Ophthalmology 2015, 122 (11), 2316–2326.

38. Li, M.; Jia, C.; Kazmierkiewicz, K. L.; Bowman, A. S.; Tian, L.; Liu, Y.; Gupta, N. A.; Gudiseva, H. V.; Yee, S. S.; Kim, M.; Dentchev, T.; Kimble, J. A.; Parker, J. S.; Messinger, J. D.; Hakonarson, H.; Curcio, C. A.; Stambolian, D., Comprehensive analysis of gene expression in human retina and supporting tissues. Human Molecular Genetics 2014, 23 (15), 4001–4014.

39. Quinn, N.; Csincsik, L.; Flynn, E.; Curcio, C.; Kiss, S.; Sadda, S.; Hogg, R.; Petö, T.; Lengyel, I., The clinical relevance of visualising the peripheral retina. Progress in Retinal and Eye Research 2018.

40. Thoreson, W. B.; Dacey, D. M., Diverse Cell Types, Circuits, and Mechanisms for Color Vision in the Vertebrate Retina. Physiological Reviews 2019, 99 (3), 1527–1573.

41. Ablonczy, Z.; Smith, N.; Anderson, D. M.; Grey, A. C.; Spraggins, J.; Koutalos, Y.; Schey, K. L.; Crouch, R. K., The utilization of fluorescence to identify the components of lipofuscin by imaging mass spectrometry. Proteomics 2014, 14 (7-8), 936–44.

42. Snodderly, D. M.; Auran, J. D.; Delori, F. C., The macular pigment. II. Spatial distribution in primate retinas. Investigative Ophthalmology & Visual Science 1984, 25 (6), 674–685.

43. Powner, M. B.; Gillies, M. C.; Tretiach, M.; Scott, A.; Guymer, R. H.; Hageman, G. S.; Fruttiger, M., Perifoveal Müller Cell Depletion in a Case of Macular Telangiectasia Type 2. Ophthalmology 2010, 117 (12), 2407–2416.

44. Powner, M. B.; Gillies, M. C.; Zhu, M.; Vevis, K.; Hunyor, A. P.; Fruttiger, M., Loss of Müller’s Cells and Photoreceptors in Macular Telangiectasia Type 2. Ophthalmology 2013, 120 (11), 2344–2352.

45. Obana, A.; Sasano, H.; Okazaki, S.; Otsuki, Y.; Seto, T.; Gohto, Y., Evidence of Carotenoid in Surgically Removed Lamellar Hole-Associated Epiretinal Proliferation. Investigative Ophthalmology & Visual Science 2017, 58 (12), 5157–5163.

46. Pang, C. E.; Maberley, D. A.; Freund, K. B.; White, V. A.; Rasmussen, S.; To, E.; Matsubara, J. A., Lamellar Hole-Associated Epiretinal Proliferation: A Clinicopathologic Correlation. RETINA 2016, 36 (7), 1408–1412.

47. Bazan, N. G., Cell survival matters: docosahexaenoic acid signaling, neuroprotection and photoreceptors. Trends in Neurosciences 2006, 29 (5), 263–271.

48. Bazan, N. G., Neuroprotectin D1 (NPD1): A DHA-Derived Mediator that Protects Brain and Retina Against Cell Injury-Induced Oxidative Stress. Brain Pathology 2005, 15 (2), 159–166.

49. Zhang, X.; Hughes, B. A., KCNQ and KCNE potassium channel subunit expression in bovine retinal pigment epithelium. Experimental Eye Research 2013, 116, 424–432.

50. Nawrot, M.; West, K.; Huang, J.; Possin, D. E.; Bretscher, A.; Crabb, J. W.; Saari, J. C., Cellular Retinaldehyde-Binding Protein Interacts with ERM-Binding Phosphoprotein 50 in Retinal Pigment Epithelium. Investigative Ophthalmology & Visual Science 2004, 45 (2), 393–401.

51. Messias, M. C. F.; Mecatti, G. C.; Priolli, D. G.; de Oliveira Carvalho, P., Plasmalogen lipids: functional mechanism and their involvement in gastrointestinal cancer. Lipids in Health and Disease 2018, 17 (1), 41.

52. Braverman, N. E.; Moser, A. B., Functions of plasmalogen lipids in health and disease. Biochimica et Biophysica Acta (BBA) - Molecular Basis of Disease 2012, 1822 (9), 1442–1452.

53. Winkler, B. S.; Boulton, M. E.; Gottsch, J. D.; Sternberg, P., Oxidative damage and age-related macular degeneration. Molecular vision 1999, 5, 32–32.

54. Jarrett, S. G.; Boulton, M. E., Consequences of oxidative stress in age-related macular degeneration. Molecular aspects of medicine 2012, 33 (4), 399–417.

55. Farooqui, A. A.; Horrocks, L. A., Book Review: Plasmalogens: Workhorse Lipids of Membranes in Normal and Injured Neurons and Glia. The Neuroscientist 2001, 7 (3), 232–245.

56. Kooijman, A. C., Light distribution on the retina of a wide-angle theoretical eye. Journal of the Optical Society of America 1983, 73 (11), 1544–1550.

57. Weale, R. A., Retinal illumination using a wide-angle model of the eye:comment. Journal of the Optical Society of America A 1990, 7 (1), 170–171.

58. Tang, P. H.; Kono, M.; Koutalos, Y.; Ablonczy, Z.; Crouch, R. K., New insights into retinoid metabolism and cycling within the retina. Progress in Retinal and Eye Research 2013, 32, 48–63.

59. Goldman, D., Müller glial cell reprogramming and retina regeneration. Nature reviews. Neuroscience 2014, 15 (7), 431–442.

60. Kiser, P. D.; Golczak, M.; Palczewski, K., Chemistry of the retinoid (visual) cycle. Chemical reviews 2014, 114 (1), 194–232.

61. Hughes, A., The Topography of Vision in Mammals of Contrasting Life Style: Comparative Optics and Retinal Organisation. In The Visual System in Vertebrates, Crescitelli, F.; Dvorak, C. A.; Eder, D. J.; Granda, A. M.; Hamasaki, D.; Holmberg, K.; Hughes, A.; Locket, N. A.; McFarland, W. N.; Meyer, D. B.; Muntz, W. R. A.; Munz, F. W.; Olson, E. C.; Reyer, R. W.; Crescitelli, F., Eds. Springer Berlin Heidelberg: Berlin, Heidelberg, 1977; pp 613–756.

