## Supplemental Figures and Table for "Lipid Landscape of the Human Retina and Supporting Tissues Revealed by High Resolution Imaging Mass Spectrometry"

**Supplement Information**

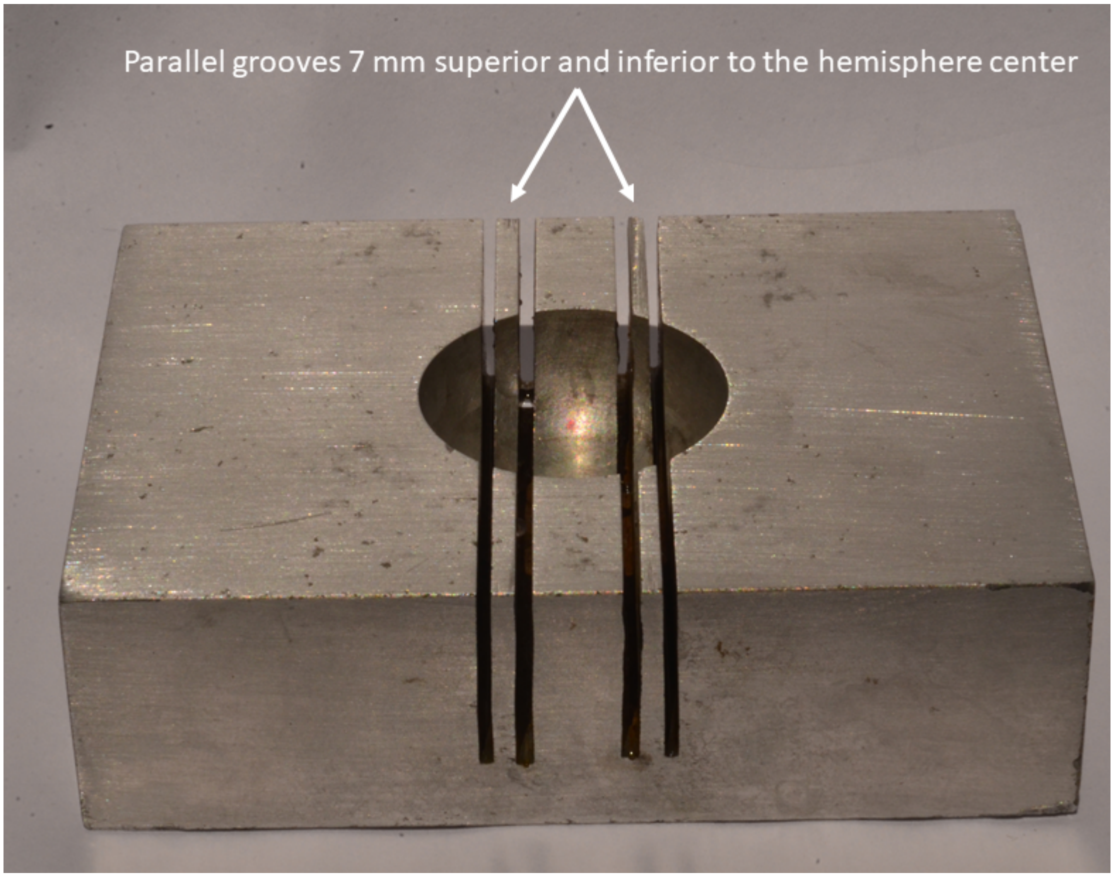


**Supplemental Figure 1. Custom-designed aluminum 3" x 4" x 1" billet for standardized dissection of a human eye.** A human eye is placed within the 30-mm-diameter hemispheric well, looking up. The parallel grooves 7 mm superior and inferior to the hemisphere center provide orientation of a tissue blade while a tissue belt through the middle of the globe is sliced out.

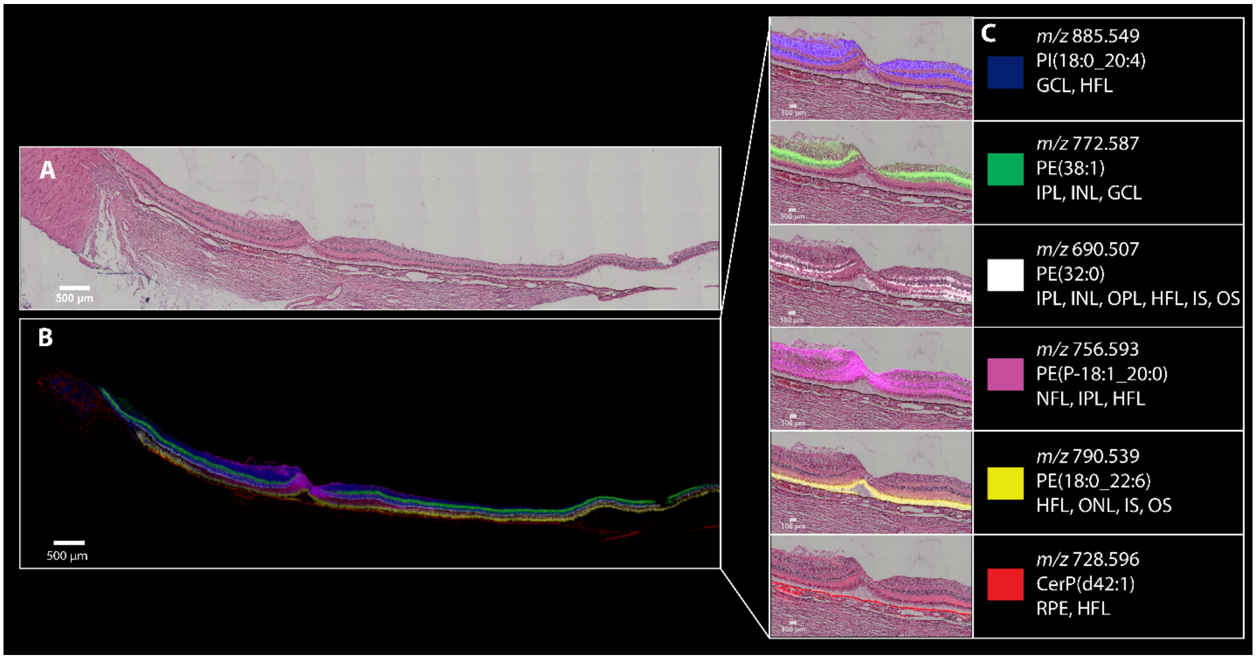


**Supplemental Figure 2. Biologic replicate of data in Figure 2 showing 6 distinct signals in human macula.** H&E and MALDI IMS images of ocular tissue from an 81-year-old donor displaying six signals with distinct cellular localizations. **A)** Un-zoomed H&E stained tissue. **B)** Overlay of MALDI IMS images showing selected lipid signals that localize to distinct layers of the neural retina (10 µm pixel size). **C)** Individual ion signals overlaid on top of H&E stained tissue image.

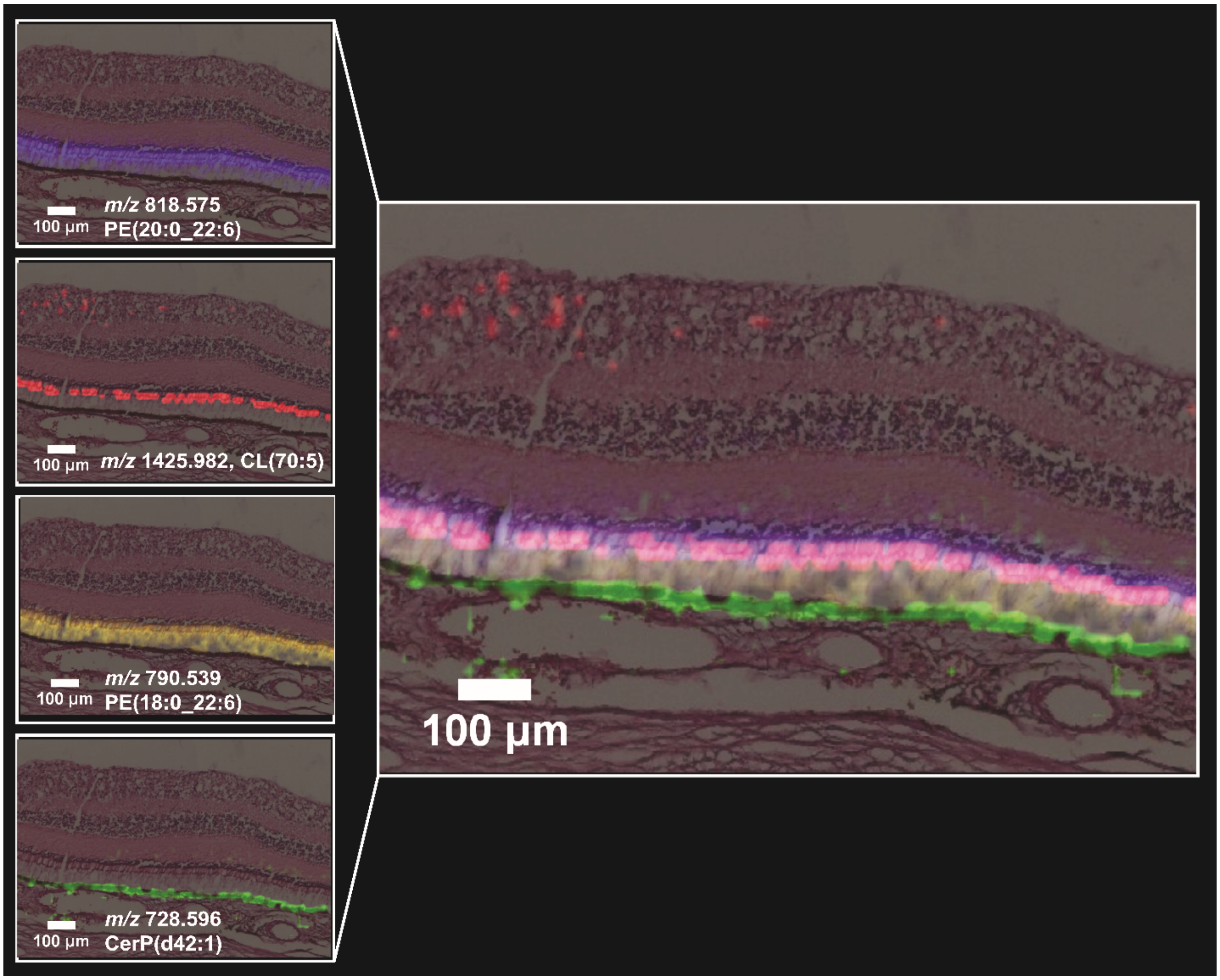


**Supplemental Figure 3. Biologic replicate of imaging mass spectrometry signals in photoreceptors shown in Figure 3.** MALDI IMS images and H&E stained tissue images from 81 year old donor overlaid in peripheral retina displaying signals that localize to subcellular compartments of outer retinal cells (15 µm pixel size). Unlike the images in Figure 3 and Supplemental 5, and like those in Figure 2, the intense signal for *m/z* 728.596 (green) localizes to a thick layer possibly representing localization in cell bodies of RPE in addition to apical processes. These differences in localization may be due to the attachment of RPE to photoreceptors, the lower resolution of scanning in this figure, or both.

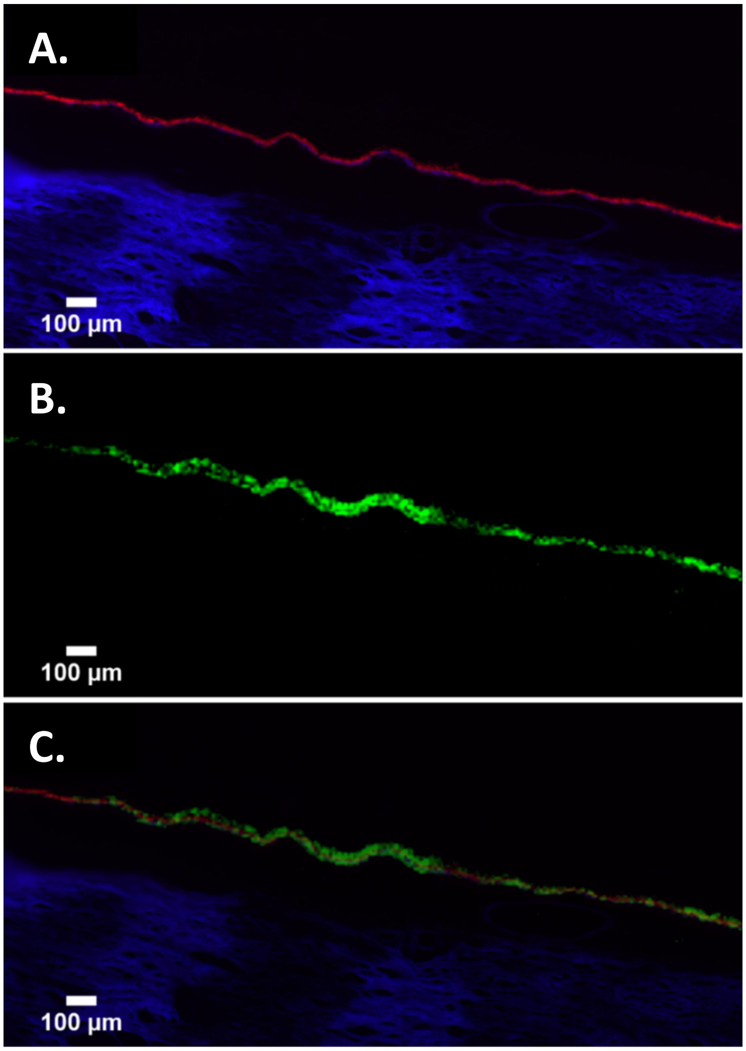


**Supplemental Figure 4. Microscopy assists imaging mass spectrometry localization. A)** Autofluorescence (AF) imaging of tissue from a 83 year old donor displaying intense signal originating from RPE (red) and sclera (blue). The choroid (between RPE and sclera) is not visible. **B)** MALDI IMS of *m/z* 728.596 (10 µm pixel size). **C)** AF overlaid with *m/z* 728.596 in green.

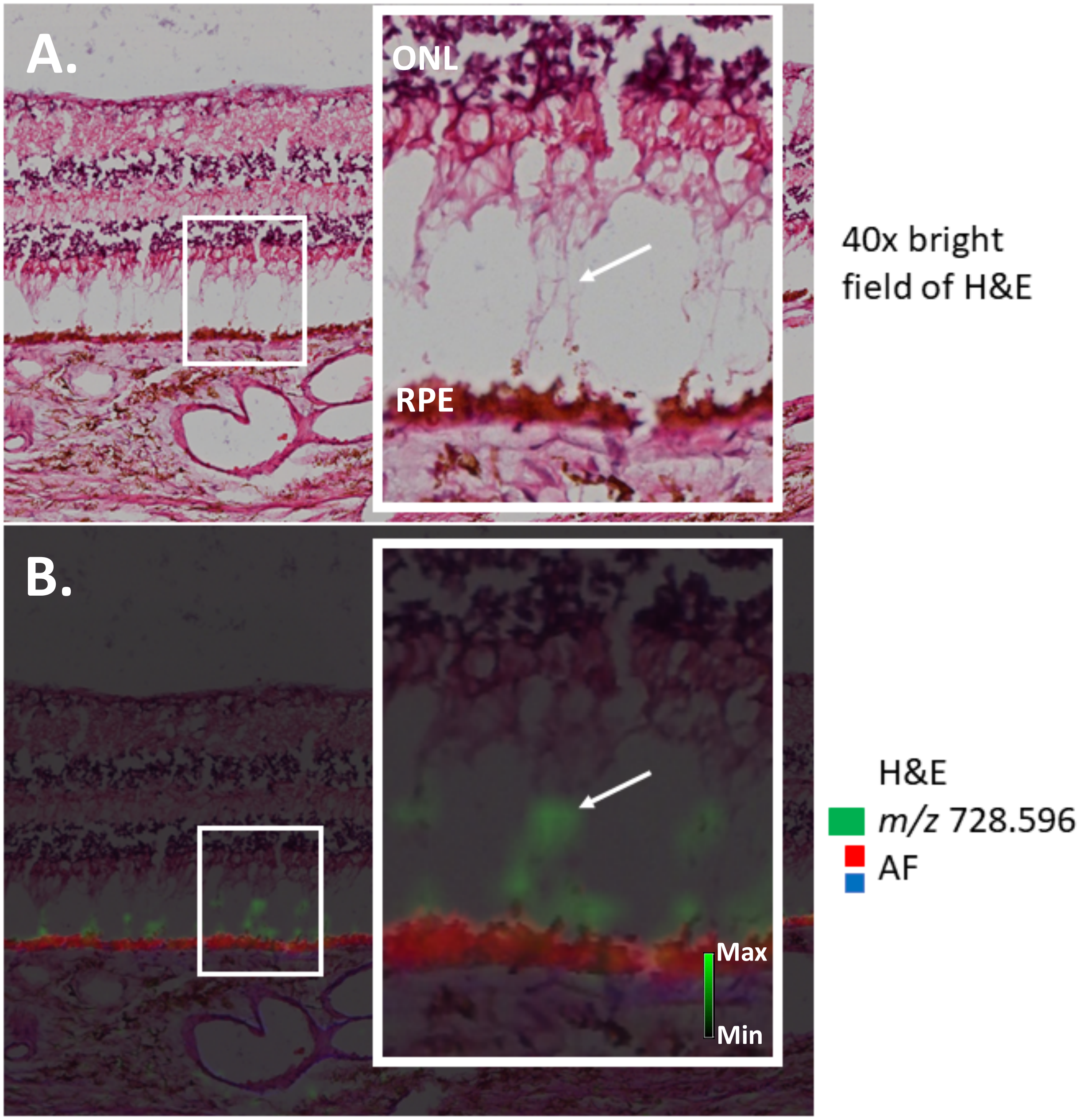


**Supplemental Figure 5. Evidence for molecular localization to RPE apical processes. A)** A 40X H&E stained tissue displaying a region where the photoreceptors have separated artifactually from the RPE. **B)** 10x H&E stained tissue overlaid with AF and MALDI IMS of *m/z* 728.596 CerP(d42:1), captured at 10 µm resolution. Image insets include arrows indicating a tissue region where this signal is present outside the RPE cell bodies, which can be seen in the overlay image in panel B inset.

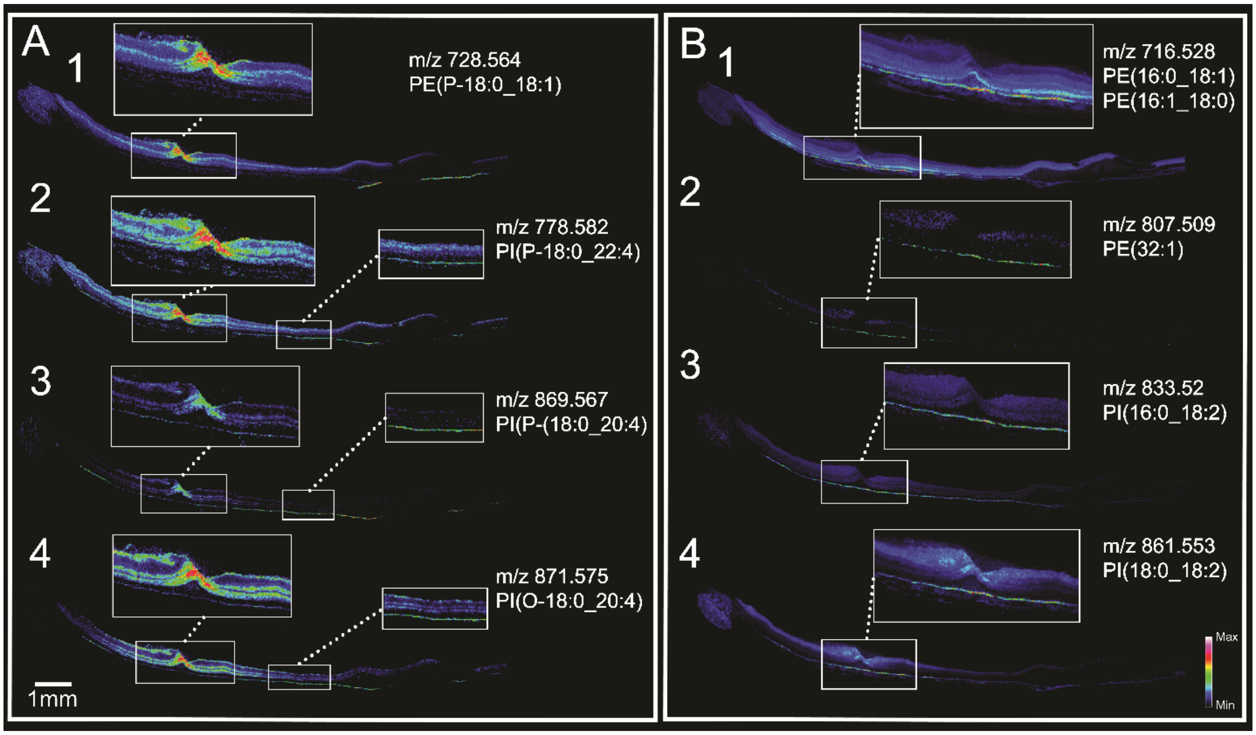


**Supplemental Figure 6. MALDI IMS images of complementary macular and RPE signals from an 81-year-old donor (15 µm pixel size).** Biologic replicate of data shown in main Figure 4. **A)** Signals localized to layers of the macular neurosensory retina and peripheral RPE. **B)** Signals localized to RPE underlying the macula and not confined to the fovea.

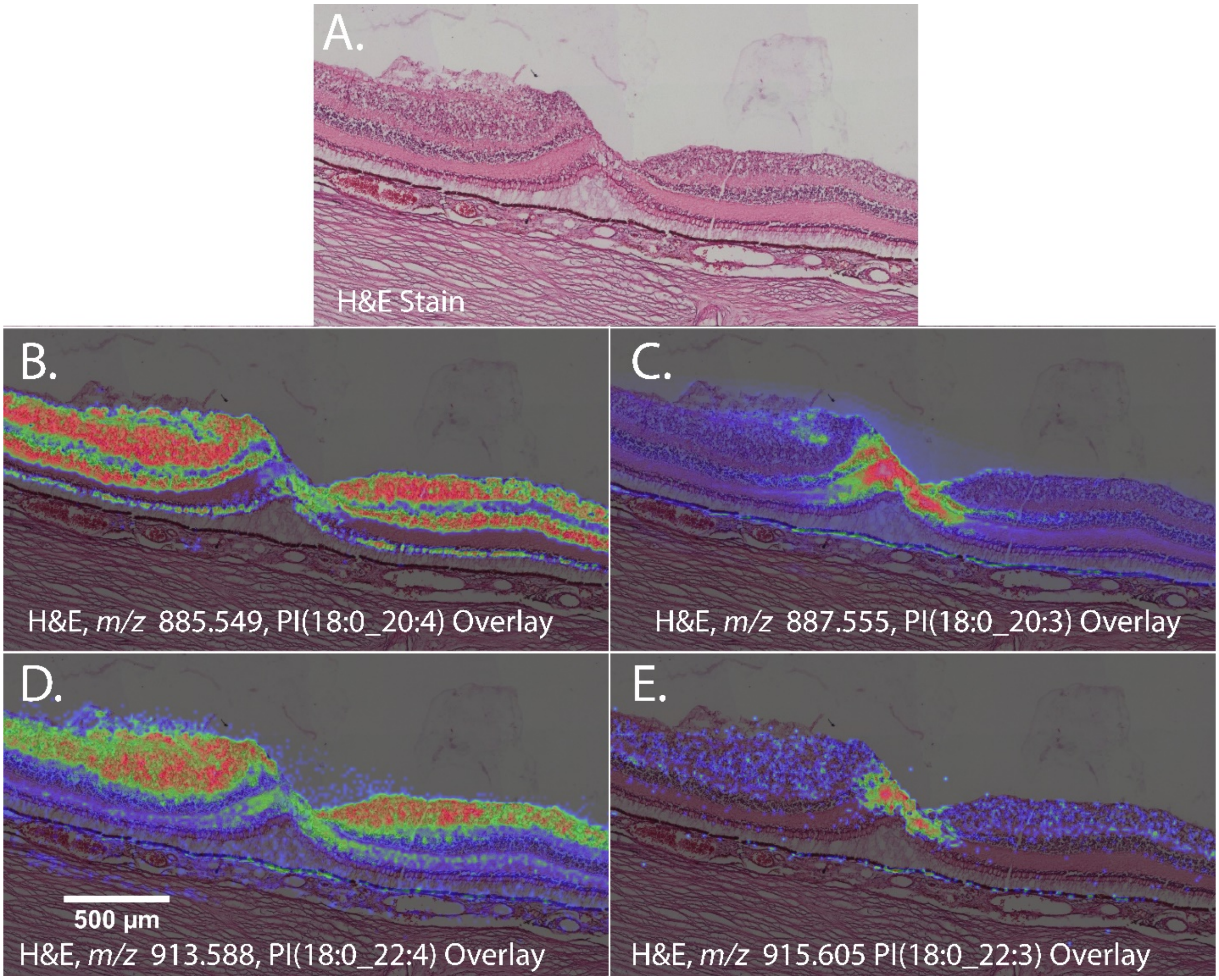


**Supplemental Figure 7. Lipid species varying in only one double bond can exhibit different MALDI IMS distributions.** Biologic replicate of data shown in main Figure 5. **A)** Zoomed H&E stained tissue image of the fovea within the macula. **B)** *m/z* 885.549, PI(18:0_20:4). **C)** *m/z* 887.555, PI(18:0_20:3). **D)** *m/z* 913.588, PI(18:0_22:4). **E)** *m/z* 915.605, PI(18:0_22:3).

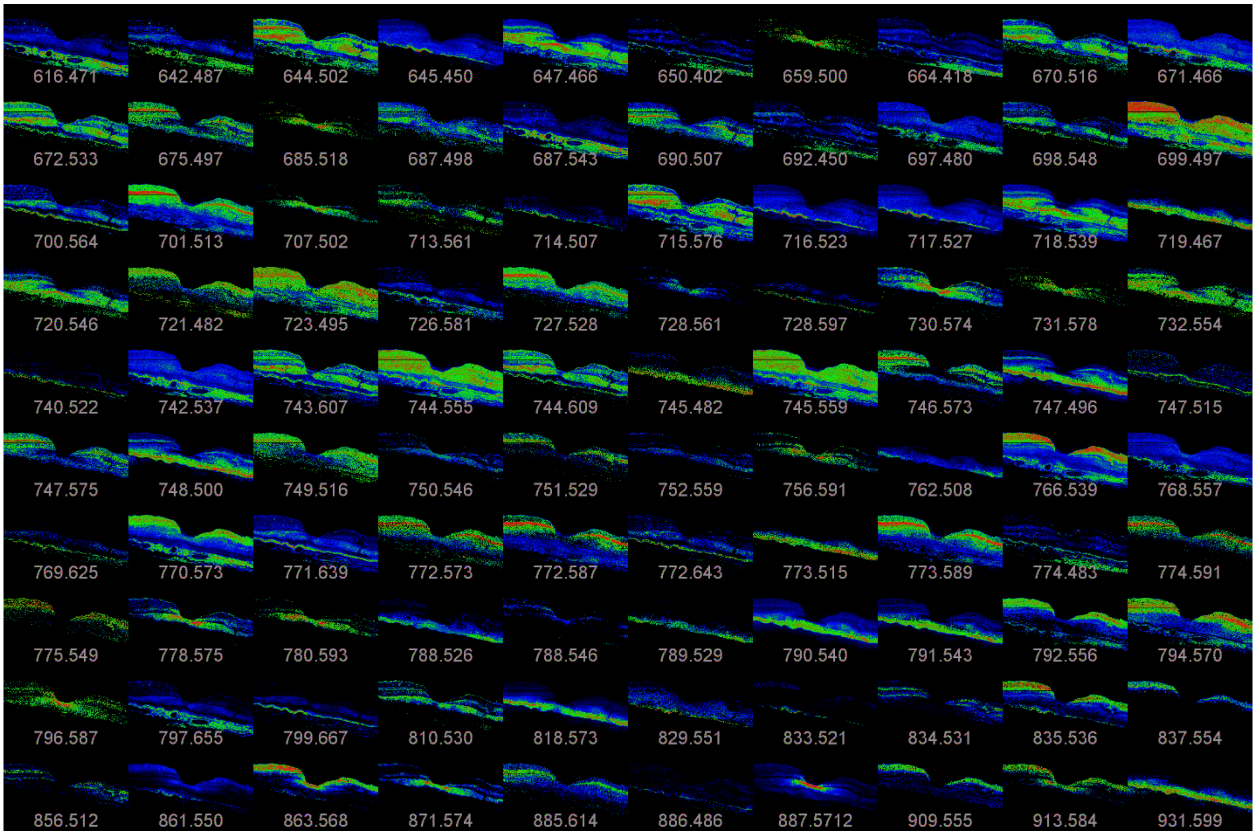


**Supplemental Figure 8. Extensive dataset obtained by imaging mass spectrometry of retina and choroid in an 83-year-old human eye donor.** From a dataset of 486 negative ion mode MALDI IMS images, 100 with qualitatively strong signals and varied localizations are shown. The macula, with foveal dip in the center, is flanked by perifovea.

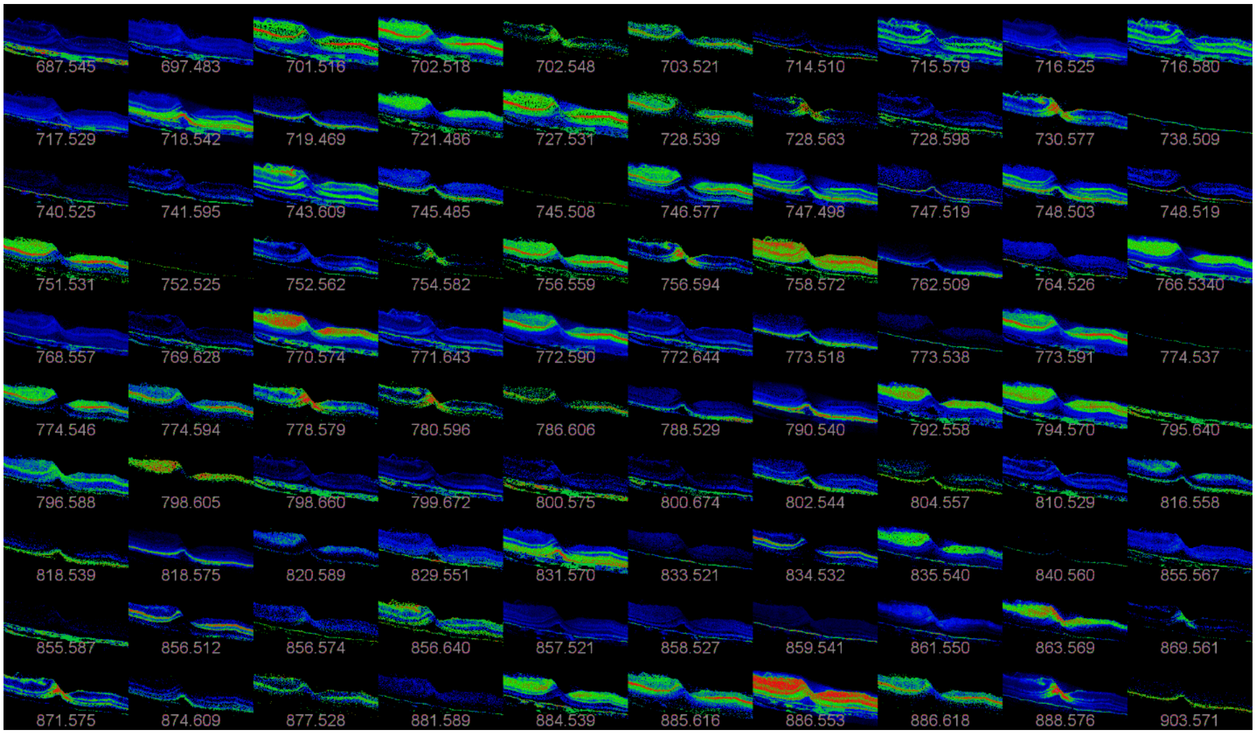


**Supplemental Figure 9. Extensive dataset obtained by imaging mass spectrometry of retina and choroid in an 81-year-old human eye donor.** From a dataset of 1,209 negative ion mode MALDI IMS images, 100 with qualitatively strong signals and varied localizations are shown. The macula, with foveal dip in the center, is flanked by perifovea.

### Supplemental Table 1. Tandem LC-MS/MS identifications from the 81-year-old donor tissue and accurate mass identification of *m/z* values from figures.

#### Tandem MS Identifications

| PE(16:0_18:1)/PE(16:1_18:0) | | *m/z* 716.526 | | |
|---|---|---|---|---|
| **m/z** | **ID** | **Th m/z** | **ppm** | **MF** |
| 253.2172 | [16:1-H]- | 253.2168 | -1.58 | C16H29O2- |
| 255.2328 | [16:0-H]- | 255.2324 | -1.57 | C16H31O2- |
| 281.2487 | [18:1-H]- | 281.2481 | -2.13 | C18H33O2- |
| 283.2655 | [18:0-H]- | 283.2637 | -6.35 | C18H35O2- |
| 478.2942 | Loss of sn2 acyl chain as ketene (RCH=C=O), [M-(16:0)-H]- | 478.2934 | -1.67 | C23H45NO7P- |
| 480.3116 | Loss of sn2 acyl chain as ketene (RCH=C=O), [M-(16:1)-H]- | 480.3091 | -5.20 | C23H47NO7P- |
| 460.2818 | [M-(16:0)-H]- | 460.2828 | 2.17 | C23H43NO6P- |
| 462.3 | [M-(16:1)-H]- | 462.2985 | -3.24 | C23H45NO6P- |
| 716.5228 | M-H- | 716.5231 | 0.42 | C39H75NO8P- |

| PE(P-18:0_18:1) | | *m/z* 728.563 | | |
|---|---|---|---|---|
| **m/z** | **ID** | **Th m/z** | **ppm** | **MF** |
| 140.0109 | Head group | 140.0114 | 3.57 | C2H7NO4P- |
| 281.2487 | [18:0-H]- | 281.2481 | -2.13 | C18H35O2- |
| 446.3026 | [M-(18:1)-H]- | 446.3036 | 2.24 | C23H45NO5P- |
| 728.5624 | M-H- | 728.56 | -3.29 | C41H79NO3P- |

| PE(P-20:0_18:1) | | *m/z* 756.593 | | |
|---|---|---|---|---|
| **m/z** | **ID** | **Th m/z** | **ppm** | **MF** |
| 281.2487 | [18:1-H]- | 281.2481 | -2.13 | C18H33O2- |
| 474.3359 | [M-(18:1)-H]- | 474.3349 | -2.11 | C25H49NO5P- |
| 492.3481 | Loss of sn2 acyl chain as ketene (RCH=C=O), [M-(18:1)-H]- | 492.3454 | -5.48 | C25H51NO6P- |
| 756.5947 | M-H- | 756.5913 | -4.49 | C43H84NO7P- |

| PE(P-18:0_22:4) | | *m/z* 778.574 | | |
|---|---|---|---|---|
| **m/z** | **ID** | **Th m/z** | **ppm** | **MF** |
| 331.2644 | [22:4-H]- | 331.2637 | -2.11 | C22H35O2- |
| 464.3144 | Loss of sn2 acyl chain as ketene (RCH=C=O), [M-(22:4)-H]- | 464.314 | -0.86 | C23H47NO6P- |
| 778.5765 | M-H- | 778.5751 | -1.80 | C45H81NO7P- |

| PE(18:0_22:6) | | *m/z* 790.539 | | |
|---|---|---|---|---|
| **m/z** | **ID** | **Th m/z** | **ppm** | **MF** |
| 283.2665 | [18:0-H]- | 283.2637 | -9.88 | C18H35O2- |
| 327.2331 | [22:6-H]- | 327.2324 | -2.14 | C22H31O2- |
| 462.2997 | [M-(22:6)-H]- | 462.2985 | -2.60 | C23H45NO6P- |
| 480.3080 | Loss of sn2 acyl chain as ketene (RCH=C=O), [M-(22:6)-H]- | 480.3091 | 2.29 | C23H47NO7P- |
| 790.5378 | M-H- | 790.5387 | 1.14 | C45H77NO8P- |

| PE(20:0/22:6) | | *m/z* 818.5748 | | |
|---|---|---|---|---|
| **m/z** | **ID** | **Th m/z** | **ppm** | **MF** |
| 327.2332 | [22:6-H]- | 327.2324 | -2.44 | C22H31O2- |
| 490.3319 | [M-(22:6)-H]- | 490.3297 | -4.49 | C25H49NO6P- |
| 508.3408 | Loss of sn2 acyl chain as ketene (RCH=C=O), [M-(22:6)-H]- | 508.3404 | -0.79 | C25H51NO7P- |
| 818.5727 | M-H- | 818.5705 | -2.69 | C47H82NO8P- |

| PI(16:0_18:2) | | *m/z* 833.523 | | |
|---|---|---|---|---|
| **m/z** | **ID** | **Th m/z** | **ppm** | **MF** |
| 255.2333 | [16:0-H]- | 255.2324 | -3.53 | C16H31O2- |
| 279.2327 | [(18:2-H]- | 279.2324 | -1.07 | C18H31O2- |
| 577.2809 | [M-(16:0)-H]- | 577.2778 | -5.37 | C27H46O11P- |
| 553.2782 | [M-(18:2)-H]- | 553.2778 | -0.72 | C25H46O11P- |
| 833.5152 | M-H- | 833.5181 | 3.48 | C43H78O13P- |

| PI(16:0_20:4) | | *m/z* 857.520 | | |
|---|---|---|---|---|
| **m/z** | **ID** | **Th m/z** | **ppm** | **MF** |
| 255.2333 | [16:0-H]- | 255.2324 | -3.53 | C16H31O2- |
| 303.2327 | [20:4-H]- | 303.2324 | -0.99 | C20H31O2- |
| 601.2759 | [M-(16:0)-H]- | 601.2778 | 3.16 | C29H46O11P- |
| 553.2786 | [M-(20:4)-H]- | 553.2778 | -1.45 | C25H46O11P- |
| 857.5206 | M-H- | 857.5181 | -2.92 | C45H78O13P- |

| PI(18:0_18:2) | | m/z 861.552 | | |
|---|---|---|---|---|
| **m/z** | **ID** | **Th m/z** | **ppm** | **MF** |
| 279.2329 | [(18:2-H]- | 279.2324 | -1.79 | C18H31O2- |
| 283.2654 | [18:0-H]- | 283.2637 | -6.00 | C18H35O2- |
| 579.2933 | [M-(18:0)-H]- | 579.2935 | 0.35 | C27H48O11P- |
| 581.3095 | [M-(18:2)-H]- | 581.3091 | -0.69 | C27H50O11P- |
| 861.5482 | M-H- | 861.5494 | 1.39 | C45H82O13P- |

| PI(P-18:0_20:4) | | *m/z* 869.555 | | |
|---|---|---|---|---|
| **m/z** | **ID** | **Th m/z** | **ppm** | **MF** |
| 241.0116 | Head group | 241.0114 | -0.83 | C6H10O8P- |
| 303.2331 | [20:4-H]- | 303.2324 | -2.31 | C20H31O2- |
| 403.2621 | [M-(20:4)-164-H]- | 403.2614 | -1.74 | C21H40O5P- |
| 565.3174 | [M-(20:4)-H]- | 565.3142 | -5.66 | C27H50O10P- |
| 869.5566 | M-H- | 869.5549 | -1.96 | C47H82O12P- |

| PI(O-18:0_20:4) | *m/z* 871.574 | | | |
|---|---|---|---|---|
| **m/z** | **ID** | **Th m/z** | **ppm** | **MF** |
| 303.233 | [20:4-H]- | 303.2324 | -1.98 | C20H31O2- |
| 405.2778 | [M-(20:4)-164-H]- | 405.277 | -1.97 | C21H42O5P- |
| 567.3306 | [M-(20:4)-H]- | 567.3299 | -1.23 | C27H52O10P- |
| 871.572 | M-H- | 871.5701 | -2.18 | C47H84O12P- |

| PI(18:0_20:4) | *m/z* 885.549 | | | |
|---|---|---|---|---|
| **m/z** | **ID** | **Th m/z** | **ppm** | **MF** |
| 283.2644 | [18:0-H]- | 283.2637 | -2.47 | C18H35O2- |
| 303.2331 | [20:4-H]- | 303.2324 | -2.31 | C20H31O2- |
| 419.2575 | eutral loss of sn2 RCOOH group and inositol from [M-164-(20:4 | 419.2562 | -3.10 | C21H40O6P- |
| 581.3107 | [M-(20:4)-H]- | 581.3091 | -2.75 | C27H50O11P- |
| 599.3188 | Loss of sn2 acyl chain as ketene (RCH=C=O), [M-(20:4)-H]- | 599.3201 | 2.17 | C27H52O12P- |
| 885.5518 | M-H- | 885.5488 | -3.39 | C47H83O13P- |

| PI(18:0_20:3) | *m/z* 887.56696 | | | |
|---|---|---|---|---|
| **m/z** | **ID** | **Th m/z** | **ppm** | **MF** |
| 241.0117 | M-H- | 241.0114 | -1.24 | C6H10O8P- |
| 283.2645 | [18:0-H]- | 283.2643 | -0.71 | C18H35O2- |
| 305.2487 | [20:3-H]- | 305.2481 | -1.97 | C20H33O2- |
| 419.2571 | Neutral loss of sn2 RCOOH group and inositol from [M-164-(2( | 419.2563 | -1.91 | C21H40O6P- |
| 581.3107 | [M-(20:3)-H]- | 581.3091 | -2.75 | C27H50O11P- |
| 887.5656 | M-H- | 887.565 | -0.68 | C47H84O13P- |

| PI(18:0_22:4) | *m/z* 913.587 | | | |
|---|---|---|---|---|
| **m/z** | **ID** | **Th m/z** | **ppm** | **MF** |
| 241.0116 | Head group | 241.0114 | -0.83 | C6H10O8P |
| 283.2643 | [18:0-H]- | 283.2643 | 0.00 | C18H35O2- |
| 331.2643 | [22:4-H]- | 331.2643 | 0.00 | C22H36O2 |
| 419.2575 | Neutral loss of sn2 RCOOH group and inositol from [M-164-(2: | 419.2563 | -2.86 | C21H40O6P- |
| 581.3099 | [M-(22:4)-H]- | 581.3091 | -1.38 | C27H50O11P- |
| 913.5828 | M-H- | 913.5812 | -1.75 | C47H82O12P- |

| PI(18:0_22:3) | *m/z* 915.601 | | | |
|---|---|---|---|---|
| **m/z** | **ID** | **Th m/z** | **ppm** | **MF** |
| 241.0115 | Head group | 241.0114 | -0.41 | C6H10O8P |
| 283.2654 | [18:0-H]- | 283.2643 | -3.88 | C18H35O2- |
| 333.2724 | [22:3-H]- | 333.2799 | 22.50 | C22H37O2- |
| 419.2577 | Neutral loss of sn2 RCOOH group and inositol from [M-164-(2: | 419.2563 | -3.34 | C21H40O6P- |
| 581.3096 | [M-(22:3)-H]- | 581.3091 | -0.86 | C27H50O11P- |
| 915.6014 | M-H- | 915.5963 | -5.57 | C49H88O13P- |

### Accurate Mass Identifications

| CerP(d42:1) | *m/z* 728.596 | | | |
|---|---|---|---|---|
| **m/z** | **ID** | **Th m/z** | **ppm** | **MF** |
| 728.596 | [CerP(d42:1)-H]- | 728.5964 | 0.55 | C42H84NO6P- |

| CL(70:5) | *m/z* 1425.9818 | | | |
|---|---|---|---|---|
| **m/z** | **ID** | **Th m/z** | **ppm** | **MF** |
| 1425.9818 | [CL(70:5)-H]- | 1425.9801 | -1.19 | C79H143O17P2- |

| PE(38:1) | *m/z* 772.5874 | | | |
|---|---|---|---|---|
| **m/z** | **ID** | **Th m/z** | **ppm** | **MF** |
| 772.5874 | [PE(38:1)-H]- | 772.5862 | -1.55 | C43H84NO8P- |

| PI(32:1) | *m/z* 807.501 | | | |
|---|---|---|---|---|
| **m/z** | **ID** | **Th m/z** | **ppm** | **MF** |
| 807.501 | [PI(32:1)-H]- | 807.5024 | 1.73 | C41H76O13P- |

| PE(32:0) | *m/z* 690.507 | | | |
|---|---|---|---|---|
| **m/z** | **ID** | **Th m/z** | **ppm** | **MF** |
| 690.507 | [PE(32:0)-H]- | 690.5074 | 0.58 | C37H73NO8P- |
